# Thermodynamics of unicellular life: Entropy production rate as function of the balanced growth rate

**DOI:** 10.1101/2024.07.13.603363

**Authors:** Maarten J. Droste, Maaike Remeijer, Robert Planqué, Frank J. Bruggeman

## Abstract

In isothermal chemical reaction networks, reaction rates depend solely on the reactant concentrations setting their thermodynamic driving force. Living cells can, in addition, alter reaction rates in their enzyme-catalysed networks by changing enzyme concentrations. This gives them control over their metabolic activities, as function of conditions. Thermodynamics dictates that the steady-state entropy production rate (EPR) of an isothermal chemical reaction network rises with its reaction rates. Here we ask whether microbial cells that change their metabolism as function of growth rate can break this relation by shifting to a metabolism with a lower thermodynamic driving force at faster growth.

We address this problem by focussing on balanced microbial growth in chemostats. Since the driving force can then be determined and the growth rate can be set, chemostats allow for the calculation of the (specific) EPR.

First we prove that the EPR of a steady-state chemical reaction network rises with its driving force. Next, we study an example metabolic network with enzyme-catalysed reactions to illustrate that maximisation of specific flux can indeed lead to selection of a pathway with a lower driving force.

Following this idea, we investigate microbes that change their metabolic network responsible for catabolism from an energetically-efficient mode to a less efficient mode as function of their growth rate. This happens for instance during a shift from complete degradation of glucose at slow growth to partial degradation at fast growth. If partial degradation liberates less free energy, fast growth can occur at a reduced driving force and possibly a reduced EPR. We analyse these metabolic shifts using three models for chemostat cultivation of the yeast *Saccharomyces cerevisiae* that are calibrated with experimental data. We also derive a criterion to predict when EPR drops after a metabolic switch that generalises to other organisms. Both analyses gave however inconclusive results, as current experimental evidence proved insufficient. We indicate which experiments are required to get a better understanding of the behaviour of the EPR during metabolic shifts in unicellular organisms.

## 1. Introduction

Living systems such as growing cells and ecosystems are considered ‘open’ thermodynamic systems [Kondepudi and Prigogine, 2014]. They maintain displaced from thermodynamic equilibrium by a continuous exchange of mass and energy with their environment and attain non-equilibrium steady-states (NESSs) in constant conditions. NESSs are characterised by a net entropy production and persist as long as the system’s driving force is not depleted. Entropy production prevents systems from reaching their maximal entropy state, thermodynamic equilibrium, which is reached when the driving force has run out. Out-of-equilibrium open systems therefore effectively feed on ‘negentropy’ [Schrödinger, 1944, Von Stockar and Liu, 1999], when they produce entropy.

Entropy production rate (EPR) has been investigated to characterise biological systems. As living organisms are also physical systems, it can be used to understand their general thermodynamic properties [Qian and Beard, 2005, Beard et al., 2004]. Next to the role of maximisation of EPR in physical systems [Martyushev and Seleznev, 2006], such a maximisation principle has critically been analysed in regard to evolution of biological complexity [Jennings et al., 2020] and ecosystems functioning [Meysman and Bruers, 2010]. Furthermore, EPR and accumulation of entropy are studied in relation to aging and blood flow rate [Balmer, 1982, Yildiz et al., 2020].

Balanced microbial growth [Schaechter, 2015], a state in which a culture of cells grows at a fixed rate and cellular metabolism is in steady state, is an example of a NESS [Bruggeman et al., 2020, Qian and Beard, 2005]. It can, for instance, be attained in a chemostat [Novick and Szilard, 1950].

Many concepts in microbial physiology derive from biological thermodynamics. Modes of growth are, for instance, distinguished by the free-energy source for synthesis of energy equivalents (ATP, NAD(P)H) and precursors (e.g., amino acids, nucleic acids) for biosynthesis of macromolecular constituents (proteins, D/RNA, etc.) [Neidhardt et al., 1990]. Novel modes of catabolism have even been discovered from the existence of net driving forces in nature [Broda, 1977] (e.g., anaerobic ammonium oxidation [Jetten et al., 1998]), as captured by the famous statement of Baas Becking: “The environment selects” (with environment interpreted as the ‘driving force’) [O’Malley, 2008].

Thermodynamics has been used to analyse the efficiency of energy transduction from catabolism to anabolism [Westerhoff et al., 1982] and offers, in principle, a method to do so without having to consider reaction kinetics, which are generally poorly known. The “energy-converter” model has in particular received ample attention [Westerhoff et al., 1982, Stucki, 1980]. Most of its insights are, however, only applicable close to thermodynamic equilibrium, when reaction rates depend linearly on their driving force, and the entropy production is minimal [Kondepudi and Prigogine, 2014]. This does occur in microbial physiology [Jackson and McInerney, 2002], but is generally not the case. Non-equilibrium extensions of this model such as mosaic non-equilibrium thermodynamics [Westerhoff et al., 1982] did allow for appreciation of a broader class of systems, but remain based on a linearisation of reaction rates (flow) with respect to their (driving) force. Key insights from these approaches are that we can predict the biomass yield from the reduction degree of the energy source reasonably well [Heijnen and Dijken, 1992, Heijnen, 1994], and that the thermodynamic efficiency of living cells is low (presumably) to maintain them in states of high growth rates [Westerhoff et al., 1983].

The development of metabolic network reconstruction methods from sequenced genomes [Francke et al., 2005] has allowed construction of genome-scale stoichiometric models (GEMs) of metabolism [Fang et al., 2020]. GEMs in principle contain all metabolic reactions encoded on a genome. Their analysis with, for instance, flux balance analysis (FBA) [Orth et al., 2010], allows for the incorporation of thermodynamic constraints [Henry et al., 2007]. FBA can predict the maximal biomass yield on the carbon source (often also the energy source). Since the molar requirement of ATP and chemical elements for 1 gram biomass are generally constant across conditions [Stouthamer, 1973], the maximal biomass yield is attained by the mode of catabolism that generates most ATP per mole of energy source. A prominent example is the complete combustion of glucose into carbon dioxide and water via aerobic respiration by *S. cerevisiae* and *E. coli*.

At high growth rates in a glucose-limited chemostat, *S. cerevisiae* and *E. coli* gradually shift metabolism from pure respiration towards overflow metabolism [Bruggeman et al., 2020, Elsemman et al., 2022, van Hoek et al., 1998, Basan et al., 2015]. Overflow metabolism involves the partial degradation of glucose into ethanol or acetate and yields less ATP per glucose than respiration. Overflow metabolism can be predicted by FBA when specific flux constraints are added [De Groot et al., 2020]. These additional constraints are nowadays rationalised by the fact that protein concentrations inside cells are limited [Basan et al., 2015, Molenaar et al., 2009]. Since faster growing cells require higher protein concentrations in anabolism, less protein remains for catabolism. Because overflow metabolism relies on fewer reactions and partially degrades glucose, it can above a critical growth rate reach a higher ATP synthesis rate per unit enzyme than respiration. As a result, cells gradually increase overflow metabolism at the expense of respiration [Elsemman et al., 2022].

The entropy production rate (EPR) is an important quantity for chemical reaction networks [Ge and Qian, 2010, Rao and Esposito, 2016], and has more recently been used to predict microbial physiology. It has been suggested to be a driver of evolution, with microbes striving for maximal EPR Unrean and Srienc [2011], and that an upper limit on Gibbs energy dissipation rate (which is directly tied to the EPR) directly influences microbial metabolism and growth Niebel et al. [2019].

Since entropy production maintains cells displaced from equilibrium, we intuitively expect that higher growth rates coincide with increased EPR (given by the product of the growth rate and thermodynamic driving force). However, entropy production is in part determined by the driving force. Can the EPR of a cell decrease as function of its growth rate when it shifts its biochemical reaction network from complete to partial glucose degradation and a reduced driving force? This would be surprising as this appears not to be possible in purely chemical reaction networks, which do not involve enzyme catalysts.

We show that the chemostat is a suitable device to study these problems, as the net driving force and rate of growth can be independently determined, allowing for a calculation of the (specific) EPR. Using a combination of stoichiometric and chemostat models, we explore the possibility that EPR does not increase with growth rate after a metabolic switch, and derive a sufficient condition for the EPR to decrease after such a switch. These results indicate that despite feeding on negentropy to stay away from thermodynamic equilibrium, higher growth rates do not necessarily involve higher entropy production rates.

## 2. Theoretical Preliminaries

### 2.1 Reaction kinetics and stoichiometric network theory

In this section, we first briefly summarise two pieces of existing theory and next we integrate them. We relate theory about (bio)chemical reactions, their stoichiometry, kinetics and thermodynamics [Qian and Beard, 2005] to theory about steady-state flux distributions of reaction networks and how they decompose into elementary flux vectors [Gagneur and Klamt, 2004]. The combined theory describes the EPR of a reaction network in terms of its stoichiometric structure and kinetics. This will enable us to contrast chemical reaction networks that lack catalytic enzymes with biochemical networks that do have them.

### 2.2 Chemical versus biochemical reaction networks

Chemical reactions differ from enzyme-catalysed biochemical reactions. Chemical reactions occur spontaneously without catalytic agents; the rate of a biochemical reaction, at constant temperature and pressure, depends on the concentration of the reactants, setting the reaction’s thermodynamic driving force, and the concentration of the catalysing enzymes and (allosteric) effectors [Cornish-Bowden, 2014]. (The enzyme is a catalyst because it does not change during the conversion; it is unaltered by the catalytic events.) A chemical reaction depends only on the driving force, set by reactant concentrations. In the absence of an enzyme, a biochemical reaction reduces to a chemical reaction and then generally proceeds much slower—which is one of the reasons why cells exploit enzymes [Cornish-Bowden, 2014].

The stoichiometry of any (bio)chemical reaction can be written as

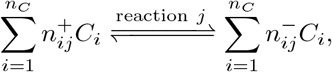

with *C*_*i*_ as the name of chemical compound *i, n*_*C*_ as the number of reactants, 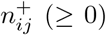 as the substrate stoichiometric constant of *C*_*i*_ in reaction *j*, 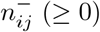 as the product stoichiometric constant of *C*_*i*_ in reaction *j*. The net stoichiometric coefficient of reactant *i* in reaction *j* equals 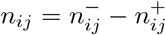; this coefficient enters in the stoichiometric matrix. The net stoichiometric coefficient for an enzyme is zero, for a substrate it is negative and for a product it is positive.

In well-mixed conditions, the rate *v*_*j*_ of chemical reaction *j* obeys mass-action kinetics

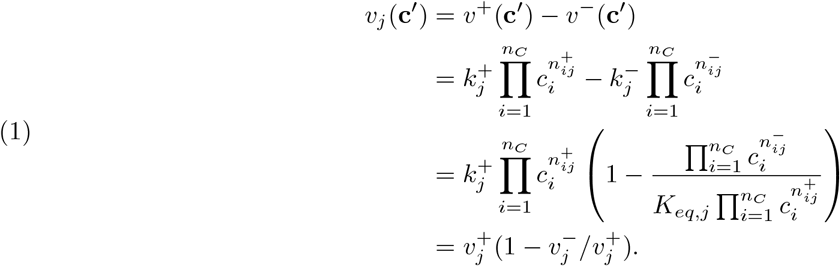

The concentration of *C*_*i*_ is denoted here as *c*_*i*_, 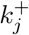 and 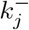 are, respectively, the forward and the backward rate constant, and *K*_*eq,j*_ is the equilibrium constant of reaction *j*. The vector **c**^*′*^ is the vector **c**, containing the concentrations of the time-dependent reactants, augmented with the concentrations of the reactants that are held fixed (they are thus seen as ‘external’ to the (bio)chemical reaction network).

The rate *v*_*j*_ of an enzyme-catalysed biochemical reaction *j* can generally be written as [Bruggeman et al., 2020],

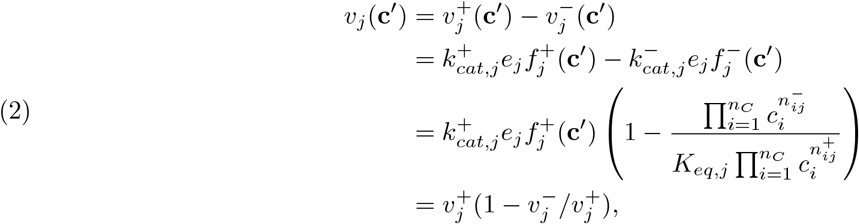

with 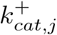 and 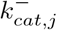 as the forward and backward catalytic rate constants, 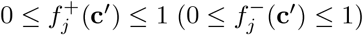 as the substrate (product) saturation function (which can contain activating or inhibiting (allosteric) effector influences, enzyme cooperativity, etc.) of enzyme *j* catalysing reaction *j*. Note that the rate of an enzymatic reaction depends on the concentration of reactants and effectors (in **c**^*′*^) and on the enzyme concentration. Effector and enzyme concentrations do not play a role in chemical reaction networks.

The term that occurs in both chemical and biochemical reaction kinetics,

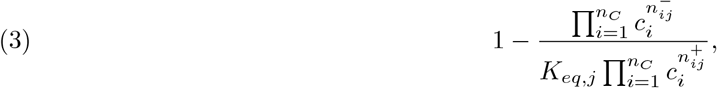

is the displacement from thermodynamic equilibrium by reaction *j*. Below we show that this term relates to the thermodynamic driving force of the reaction.

The stoichiometric matrix of a chemical reaction network **N** with entries {**N**}_*ij*_ = *n*_*ij*_ contains all the net stoichiometric coefficients of the reactants with variable (i.e., time-dependent) concentrations.

The rate of change in the concentration of the reactants is now given by

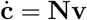

and steady state implies

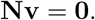

### 2.3 Reaction thermodynamics

The (molar) Gibbs free energy potential of reaction *j* equals,

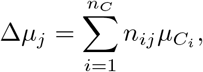

where 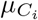 is the chemical potential or (partial) Gibbs free energy per mole of compound *C*_*i*_,

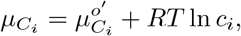

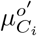 is the (partial) Gibbs free energy of formation per mole under standard conditions^1^, *R* is the ideal gas constant and *T* is the temperature.

When reaction *j* operates at thermodynamic equilibrium, Δ*μ*_*j*_ = 0, such that

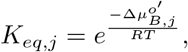

where 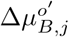 is the standard Gibbs energy of reaction under biological standard conditions, as explained in Appendix A. This leads to a relation for the rate of a (bio)chemical reaction *j* in terms of its Gibbs free energy potential Δ*μ*_*j*_, via the displacement from thermodynamic equilibrium term (3),

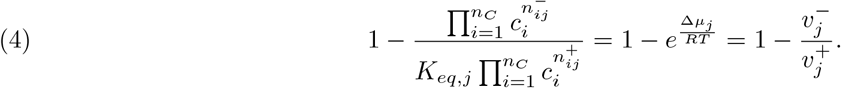

Thus at thermodynamic equilbrium: 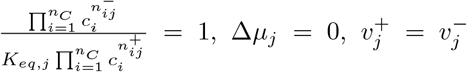 and *v*_*j*_ = 0. Note that *sign*(*v*_*j*_) = −*sign*(Δ*μ*_*j*_).

### 2.4. The rate of a (bio)chemical reaction rises with its driving force

Mass-action kinetics (1) and enzyme kinetics (2) both depend on the Gibbs free energy potential of the reaction via the term shown in eq. (3). Since the Gibbs free energy potential of a reaction is determined by the concentration of the reactants, a change in the concentration of reactants changes the free energy potential as well as the substrate saturation function in enzyme kinetics. The quantity *X*_*j*_ ≡ −Δ*μ*_*j*_ represents the thermodynamic driving force of the reaction. As *sign*(*v*_*j*_) = *sign*(*X*_*j*_), this is a more convenient quantity to consider in this paper. We note that often the driving force of a reaction system is defined as Δ*μ/T* ; here we deviate from this definition. Since the substrate saturation function of an enzyme generally rises with higher substrate and lower product concentrations and because this also increases the driving force of the reaction, the substrate saturation function generally rises with the driving force too. (Note that allosteric regulators or effectors of enzymes, which are not converted into products, and enzyme concentrations do not influence the driving force of the reaction.) Hence the rate of a (bio)chemical reaction rises with its driving force.

### 2.5. The net conversion and entropy production rate of a steady-state reaction network

We consider a reaction network at a steady state. Steady states are achieved by keeping some reactant concentrations fixed at values that lead to a net, nonzero thermodynamic driving force of the network. Each steady-state reaction network brings about a net conversion of ‘substrate’ into ‘product’ reactants. The concentrations of these compounds then eventually determine the Gibbs free energy potential of the entire chemical reaction network. The direction of the net conversion is such that the total free energy of the products is lower than that of the substrates, i.e. its driving force is positive.

Any steady-state flux vector **v** satisfies **Nv** = **0**. As illustrated in equations (1) and (2), reversible reactions may be split into two irreversible ones, and the stoichiometric matrix may be extended so that each column corresponds with either a forward or a backward reaction [Gagneur and Klamt, 2004]. We may thus assume by suitably extending **N** that all flux vectors have nonnegative entries. The set {**v** | **Nv** = **0**, *v*_*j*_ ≥ 0} forms a pointed polyhedral cone and is spanned by its extreme rays [Schrijver, 1986]: any flux vector can be written as a linear combination of extreme rays using only nonnegative coefficients. Such linear combinations are called conic combinations. In the context of metabolic pathways, the extreme rays are called Elementary Flux Modes (EFMs) [Schuster and Hilgetag, 1994]. The EFMs are uniquely determined by **N** alone.

A key property of EFMs is that each reaction in the EFM is essential to carry a steady-state flux through the entire EFM. This means that EFMs cannot be further decomposed into simpler networks that sustain a steady-state flux, and that the ratios of flux values in an EFM are constant (and determined by stoichiometry). They are ‘one-degree-of-freedom’ vectors, in the sense that one flux value determines them all. This allows for normalisation of an EFM with respect to one of its entries, a property that is useful for this work.

Denoting the steady-state flux vector of a network by **v**(**c**^*′*^), we thus have

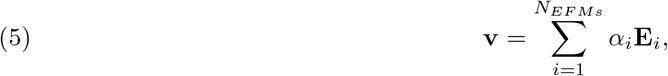

with *α*_*i*_ ≥ 0 and **E**_*i*_ as the flux vector of EFM *i* (hence, **NE**_**i**_ = 0 as well).

We introduce the stoichiometric matrix **N**^*′*^, which equals **N** augmented with the stoichiometric coefficients of the reactants that are held fixed in concentration. For each such a fixed reactant, a new row is added to **N**. The net conversion of the reaction network is referred to as the macrochemical equation (*ME*) [Heijnen and Kleerebezem, 2010]. It is used in microbial physiology, specifically in thermodynamic and yield analyses.

It can be obtained from

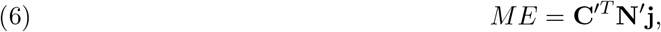

where **j** is a vector containing the number of moles of reactants consumed and/or produced (i.e. yields) in the reactions, and **C**^*′*^ is the vector with the names of the reactants, including the fixed ones.

The macrochemical equation can for instance equal *ME* = 5*R* + *S* − 3*P* − 2*Q*, which corresponds to the net reaction 3*P* + 2*Q* → 5*R* + *S* in which 3 moles of *P* and 2 moles of *Q* are converted (directly or through a reaction network) into 5 moles of *R* and one of *S*.

The Gibbs free energy potential of the steady-state chemical reaction network can also be obtained from the Gibbs free energies of the reactants [Beard et al., 2004],

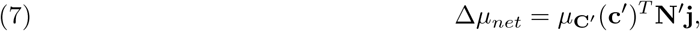

with *μ*_**C**_*′* (**c**) as the vector of the Gibbs free energies per mole of all the compounds. For the example *ME* just discussed, the Gibbs free energy potential of the (‘net’) chemical reaction network would thus be Δ*μ*_*net*_ = 5*μ*_*R*_ + *μ*_*S*_ − 3*μ*_*P*_ − 2*μ*_*Q*_. Its driving force equals *X*_*net*_ = −Δ*μ*_*net*_.

The vector **j** is closely related to the flux v ector **v**. F or a n E FM, w e m ay i n f act s ubstitute **j** b y **v**, normalised appropriately. For example, the reaction scheme

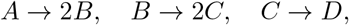

has a net reaction *A* → 4*D* and EFM equal to (1 2 4 4)^*T*^, which is uniquely determined up to a constant. Thus we take **j** = (1 2 4 4)^*T*^, so that *A* → 4*D* corresponds to Δ*μ*_*net*_ = 4*μ*_*D*_ − *μ*_*A*_.

When EFMs are conically combined as in (5) the situation is more involved. The exact conic combination depends not only on the concentration of external substrates and products, but also on the internal reaction rates. Consider the simple branched scheme

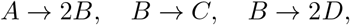

in which *B* is converted into *C* or into *D* (or both). There are two EFMs, **E**_1_ = (1 2 2 0)^*T*^ and **E**_2_ = (1 2 0 4)^*T*^, and two corresponding macrochemical equations,

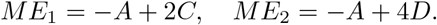

The steady state flux is a conic combination

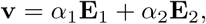

where *α*_1_ and *α*_2_ depend at least on the steady state concentrations of *A, B, C* and *D* and reaction parameters; if the reactions are additionally catalysed by enzymes, their concentration also determines the *α*_*i*_’s. We can therefore not deduce a priori the molar quantities of *C* and *D* that are produced per mole of substrate *A* consumed: these quantities depend on the *α*_*i*_. Normalising on one mole of *A* consumed, we find t hat the *ME* is

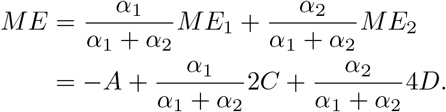

In general, provided we normalise appropriately,

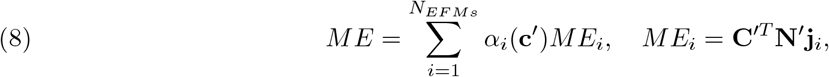

and

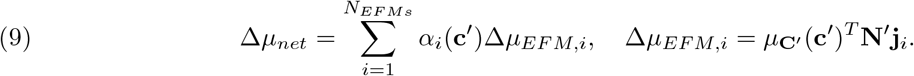

The entropy production rate (EPR) Φ of a reaction network consisting of *r* reactions is now defined as (with *S* as the entropy),

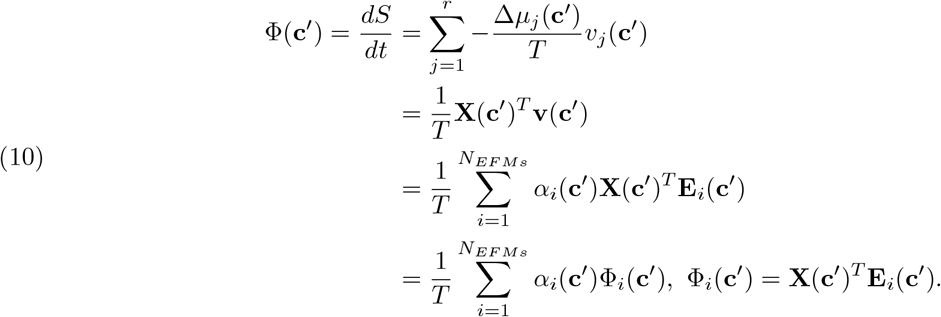

The steady-state EPR of a chemical reaction network is the weighted sum of the entropy production rates Φ_*i*_(**c**^*′*^) of its EFMs. This is the main quantity of interest in this work.

### 2.6. Modeling microbial growth and the entropy production rate in a chemostat

A chemostat is a continuously stirred bioreactor operated at a constant volume. Fresh medium with nutrients flows into the bioreactor at the same rate as medium plus biomass is removed from the bioreactor. Only when the growth rate can compensate for the loss of biomass by outflow can a microbial culture sustain itself in the bioreactor. Because inflow (not containing biomass) and outflow (containing biomass) occur at the same rate *F*, the vessel has a constant volume *V* and a dilution rate *D* = *F/V*. The growth rate *λ* of the microbe needs to equal *D* in order for a steady state to settle.

Microbial growth in the chemostat is generally modelled by condensing the process of cellular growth to the macrochemical equation (6). This reaction specifies the overall conversion of nutrients into biomass and (fermentation) products, and is included using a suitable rate equation (e.g., the Monod equation). This is rather straightforward in a chemostat, because this rate is then determined only by a single nutrient concentration called the growth-limiting substrate. The macrochemical equation can either be obtained directly from experiments, or from a flux vector that is predicted from flux balance analysis of a genomescale stoichiometric model, as in eq. (6). An example that we will be used later, see eq. (20), is

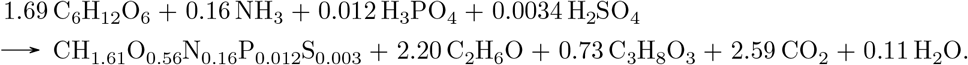

The first ‘compound’ on the right hand side of the reaction is the chemical composition of biomass, expressed per *C*-mole. The macrochemical equation is normalised to the production of 1 *C*-mole of biomass, and thus generally has the form

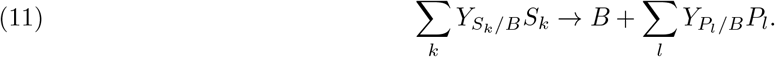

The coefficients *Y*_*i/B*_ are the yields of compounds *C*_*i*_ per *C*-mole biomass. If growth occurs at a rate *λ*, the corresponding uptake or excretion rate of compound *i* is now defined as *q*_*i/B*_ = *Y*_*i/B*_*λ*.

To model the dynamics of the chemostat we extend its well-known model description [Kuenen, 2019] with thermodynamics. The biomass concentration *b* changes due to growth and dilution as

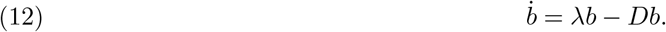

Nutrients *S*_*k*_ enter the vessel from the reservoir medium at concentrations *s*_*R,k*_ and decrease in concentration through outflow and consumption,

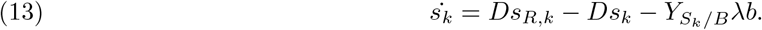

Products *P*_*l*_ are excreted by cells at rate 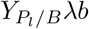 and flow out,

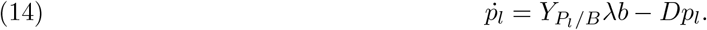

A standard implementation of the growth rate uses the Monod equation,

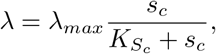

where *s*_*c*_ is the concentration of the growth-limiting carbon source 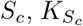 is its Monod saturation or affinity constant and *λ*_*max*_ is the maximal growth rate (in batch). We extend this model with thermodynamics by considering the Gibbs free energy potential for the macrochemical equation (11), obtained from applying (4),

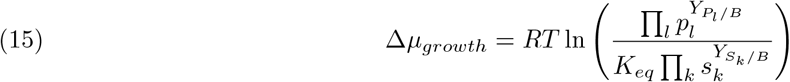

The biomass concentration *b* is not contained in the concentration term, as biomass is a solid that is not dissolved in the chemostat. The Monod equation for *λ* is now extended with the displacement from thermodynamic equilibrium,

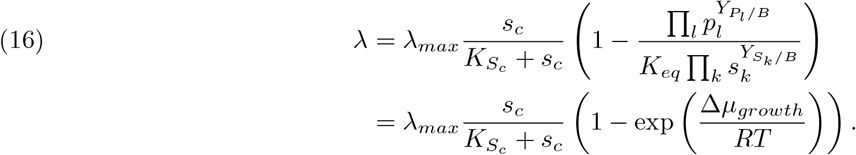

At steady state, *λ* = *D*, so the experimenter has control over the growth rate of the culture. The steady-state concentrations of substrates, products and biomass are determined by

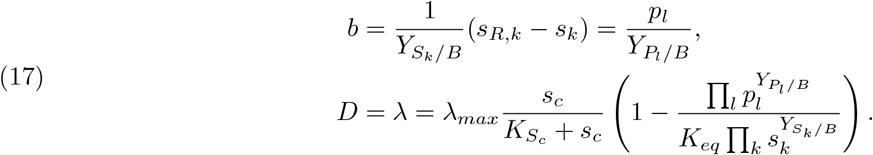

These steady states are thus determined by *D*, and we will use *b*(*D*), *s*_*k*_(*D*), etc, in appropriate places to highlight this dependence. It is easy to see that if any *s*_*k*_ decreases in steady state, then all other substrate concentrations must decrease as well, and all product concentrations must increase. These changes all affect the growth rate by lowering it. Hence, an increase in growth rate can only coincide with an increase in substrate in the medium, and a decrease in product: 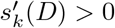 and 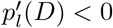. We also note that *b*^*′*^(*D*) *<* 0.

Usually the concentration of the limiting substrate in the reservoir medium is very high, i.e., 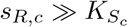.

Under this condition, the maximal dilution rate *D*_*max*_, above which wash-out of cells is faster than growth, approaches the maximal growth rate *λ*_*max*_ [Kuenen, 2019]. For microbes growing far from equilibrium, irreversibility of the macrochemical equation is approximated via a high *K*_*eq*_, i.e., a very negative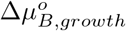.

Lastly, recall (10) for the EPR of a reaction network. Since a chemostat at steady state has fixed substrate, biomass and product concentrations, the Gibbs energy potential may be controlled, and thus compared to the growth rate. For a culture growing at a rate *λ* = *D* the (temperature-scaled) EPR of the steady-state microbial culture in the chemostat is

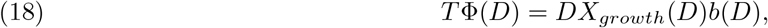

where we used that *X*_*growth*_ = −Δ*μ*_*growth*_. We are more interested in the specific (or per-capita) entropy production rate (sEPR) rather than the EPR of the entire culture. The sEPR equals

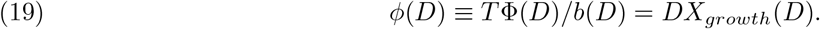

This quantity is a better measure for EPR of microbial growth in the chemostat, because the biomass concentration is also a function of growth rate. In order to analyse the EPR at different dilution rates it is therefore better to compute the EPR per unit biomass.

The chemostat model is summarised by (12)–(16). The main ingredient is a choice of macrochemical equation, which is incorporated into (16). We will explore different scenarios, in which either a culture uses a fixed macrochemical equation at different growth rates, or gradually shifts between different ones as a result of changes in protein concentrations as function of conditions. For these different scenarios we study how the sEPR (19) of the culture behaves as growth rate increases.

## 3. Results

### 3.1. Entropy production rate increases with rate in chemical reaction networks

The driving force *X* = −Δ*μ* of a chemical reaction network operating at a fixed temperature and pressure can be increased by increasing substrate concentrations or decreasing product concentrations. From the theory outlined above, we know that: i. the rate of a chemical reaction rises with its driving force according to eq. (4); ii. the driving force of a network equals a weighted sum of the driving forces of its reactions (eq. (9); it is ‘partitioned’ over the reactions); iii. the steady-state flux distribution of a network is a weighted sum of the flux distributions of its EFMs (5), and each of them is completely determined by a single flux value only. From this, it follows that a chemical reaction network with a fixed macrochemical equation (i.e., fixed conic weights for its EFMs) described by mass-action kinetics will always show an increase in EPR when the driving force of its EFMs rises, because then also all the reaction rates will rise. This can be interpreted as: processes that run at a higher rate also dissipate more energy per unit of time. The details of this argument are given in Appendix C. The same conclusion holds for a biochemical reaction network with fixed enzyme concentrations.

### 3.2. EPR may decrease with growth rate in enzyme-catalysed reaction networks

When the net driving force of a chemical reaction network with a fixed macrochemical equation is changed, all reactions that had a nonzero rate continue to do so. The relative contribution of the network’s EFMs to the steady-state flux distribution generally depends on the driving force, but EFMs do not appear or disappear by a change in the external concentration of substrates and/or products. This situation is different in biochemical reaction networks: cells can control the concentration of enzymes by gene expression, and thereby have control which reactions actually run. This control is sometimes independent of the driving force, but it may also influence it directly; by expressing genes that correspond to new reactions, new products may be excreted. The network composed out of those new EFMs could therefore have a driving force that is lower than the driving force of the set of EFMs it replaces. This raises the possibility that an increase in growth rate, for instance due to a higher nutrient concentration, may coincide with a *drop* in sEPR when cells start using new EFMs making different products.

This is particularly relevant for microbial cells that shift from an energetically-efficient metabolism at slow growth to a more inefficient mode at fast growth. Many microbes that do this show the well known shift from glucose respiration to overflow metabolism under aerobic conditions [Bruggeman et al., 2020]. The energetically-efficient mode of metabolism involves a metabolic network that, for instance, completely combusts glucose into carbon dioxide and water whereas the inefficient mode only degrades glucose partially into overflow by-products (e.g., acetate, ethanol, formate, lactate, etc.) and carbon dioxide. The inefficient mode of metabolism involves less reactions and generally extracts less free energy from glucose. So, it likely has a lower driving force, yet it is used at a higher growth rate. Such a metabolic shift could therefore be accompanied by a drop in sEPR at increasing growth rate. Below we will discuss this mechanism in the context of experimental data, but we will first show the principle with a model of a small metabolic network.

### 3.3. Maximisation of specific flux can lead to selection of a pathway with a lower driving force: a numerical illustration

Consider the metabolic network in Figure 1 composed out of seven reversible reactions, each described by enzyme kinetics, and in which the substrate *S* is consumed to produce products *P*_0_, *P*_1_ and/or *P*_2_. Concentrations of these external metabolites are considered fixed. The network has two EFMs, which can attain a steady state by themselves, or by mixing them together. Their respective driving forces are fixed at 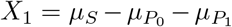 and 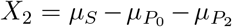. At steady state, we have the flux balances

**Figure 1.**
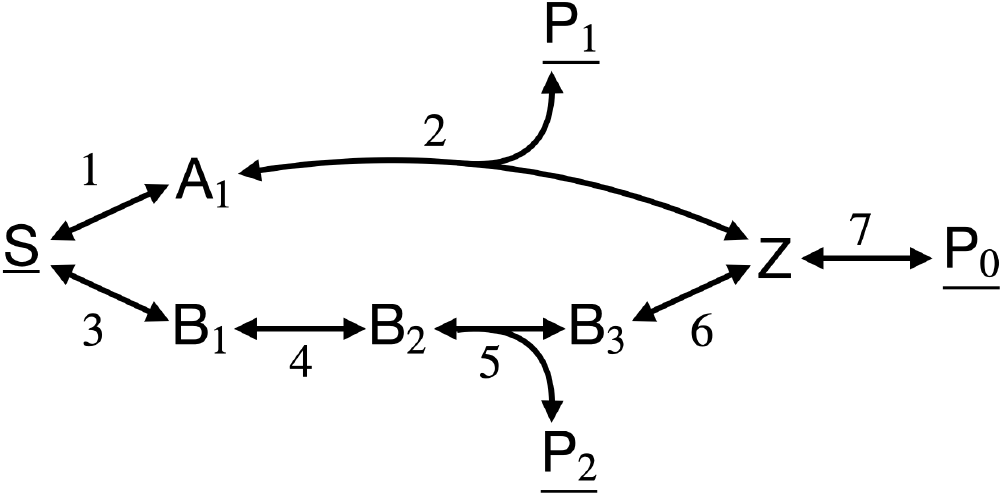
Example metabolic network to illustrate that EPR may decrease with flux. The concentration of the underlined chemical compounds are considered fixed. EFM 1 is composed out of 3 reactions (1, 2 and 7) and EFM 2 out of 5 reactions (3 to 7). EFM 1 has as thermodynamic driving force 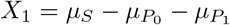 and for EFM 2 the driving force is 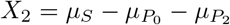.

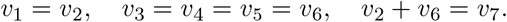

The flux through EFM 1 is taken to be *v*_1_ and denoted *J*_1_, while for EFM 2 we choose *J*_2_ = *v*_3_. The EPR of EFM 1 thus equals *X*_1_*J*_1_ and that of EFM 2 equals *X*_2_*J*_2_.

First, we consider the two EFMs separately. If the network would have been modelled as a chemical reaction network, then *J*_1_ rises with *X*_1_ and *J*_2_ rises with *X*_2_. An increase in the driving force of an EFM thus always leads to an increased EPR. Their combined EPR would then also necessarily increase.

Next, we consider the reaction network of Figure 1 as a metabolic network with the product *P*_0_ as a necessary metabolite required for cell growth, and with reaction kinetics specified by enzyme kinetic rate functions. The cell can make *P*_0_ via EFM 1 or 2. We assume that the cell chooses the pathway that gives rise to the highest steady-state synthesis flux of *P*_0_, i.e., the rate of reaction 7, per unit of total enzyme concentration invested in the metabolic network. We are therefore considering the maximisation of (*J*_1_ + *J*_2_)*/e*_*T*_ with *e*_*T*_ as the sum of the enzyme concentrations of the network. From existing theory [Wortel et al., 2014] we know that the cell will select just one of the two EFMs.

Which EFM maximises the specific flux depends on the kinetic properties of the network and the (fixed) external concentrations. Changing the external concentrations might result in a different EFM that attains the highest flux, as is illustrated in Figure 2. It shows the results of a numerical implementation of the reaction network of Figure 1. By tuning kinetic parameters, specifically catalytic rate constants *k*_*cat,j*_, the network can be parameterised such that EFM 2 maximises the flux for low concentrations of the substrate *S*. Investing the same total enzyme concentration *e*_*T*_ entirely in EFM 1 or EFM 2, the maximal value of *J*_2_ thus exceeds the maximal value of *J*_1_. As *S* increases, the flux-maximising EFM switches from EFM 2 to EFM 1. Furthermore, by setting appropriate external concentrations we can guarantee that *X*_2_ *> X*_1_ for all values of *S*. So, the switch in the optimal pathway goes together with a drop in the driving force of the network. This behaviour can not arise in the chemical reaction networks discussed in the previous section and is a direct result of the extra degrees of freedom in the form of enzyme concentrations a cell has at its disposal.

**Figure 2.**
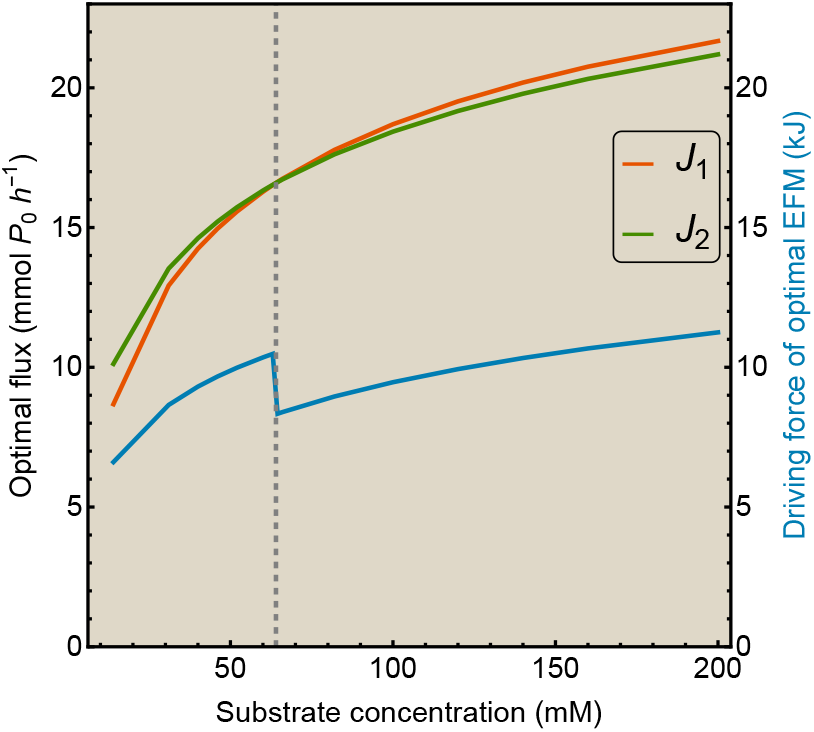
Maximisation of specific flux can lead to selection of a pathway with a lower driving force. For both EFMs 1 and 2, its respective optimal fluxes *J*_1_ and *J*_2_ are computed numerically at varying values for the concentration of *S*. At the point where *J*_1_ *> J*_2_, indicated by the gray dashed line, the corresponding driving force of the optimal EFM depicted in blue drops suddenly, as *X*_1_ *< X*_2_.

The effect of this switch on the EPR of the metabolic network can differ. We may tune the kinetic parameters such that at high *S* concentrations we obtain *J*_1_ *> J*_2_, while *X*_1_*J*_1_ *< X*_2_*J*_2_. In this case the decrease in driving force by switching from EFM 2 to EFM 1 overcompensates the increase in flux, resulting in an overall decrease of their product, i.e., the EPR. This will be analysed more thoroughly later in this paper. Nevertheless, we can already conclude that maximisation of the metabolic rate does not automatically lead to maximisation of the EPR. Generalising this to the complete metabolic network of the cell implies that EPR may decrease with growth rate in enzyme-catalysed reaction networks.

Appendix C contains general considerations for a thermodynamic consistent and feasible construction of such an example network, together with the explicit conditions for the example network presented here.

### 3.4. A constant metabolic strategy leads to a rising specific entropy production rate as function of the growth rate in the chemostat

As explained in the Theory, the chemostat provides a way of studying the (specific) EPR while controlling the growth rate. It allows for the independent measurement of the growth rate (equal to the dilution rate set by the experimenter) and the concentrations of the reactants appearing in the macrochemical equation associated with cell growth. With these ingredients, and the standard Gibbs free energies of formation of the reactants, the driving force and the sEPR can be calculated as function of the growth rate.

When the macrochemical equation is fixed, i.e., when the underlying metabolic network exploits a fixed combination of EFMs, then the driving force and the sEPR both rise with the dilution rate. For instance, when we consider the anaerobic growth of *Saccharomyces cerevisiae* on glucose (C_6_H_12_O_6_), it ferments it to ethanol (C_2_H_6_O) and glycerol (C_3_H_8_O_3_), while forming biomass (CH_1.61_O_0.56_N_0.16_P_0.012_S_0.003_), according to

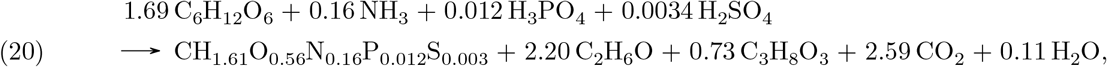

with ammonium NH_3_ as nitrogen source, H_3_PO_4_ as phosphorus source, and H_2_SO_4_ as sulphur source [Battley, 2013]. The results of the chemostat model for this macrochemical equation are depicted in Figure 3. Only concentrations of carbohydrates are shown for visual clarity.

**Figure 3.**
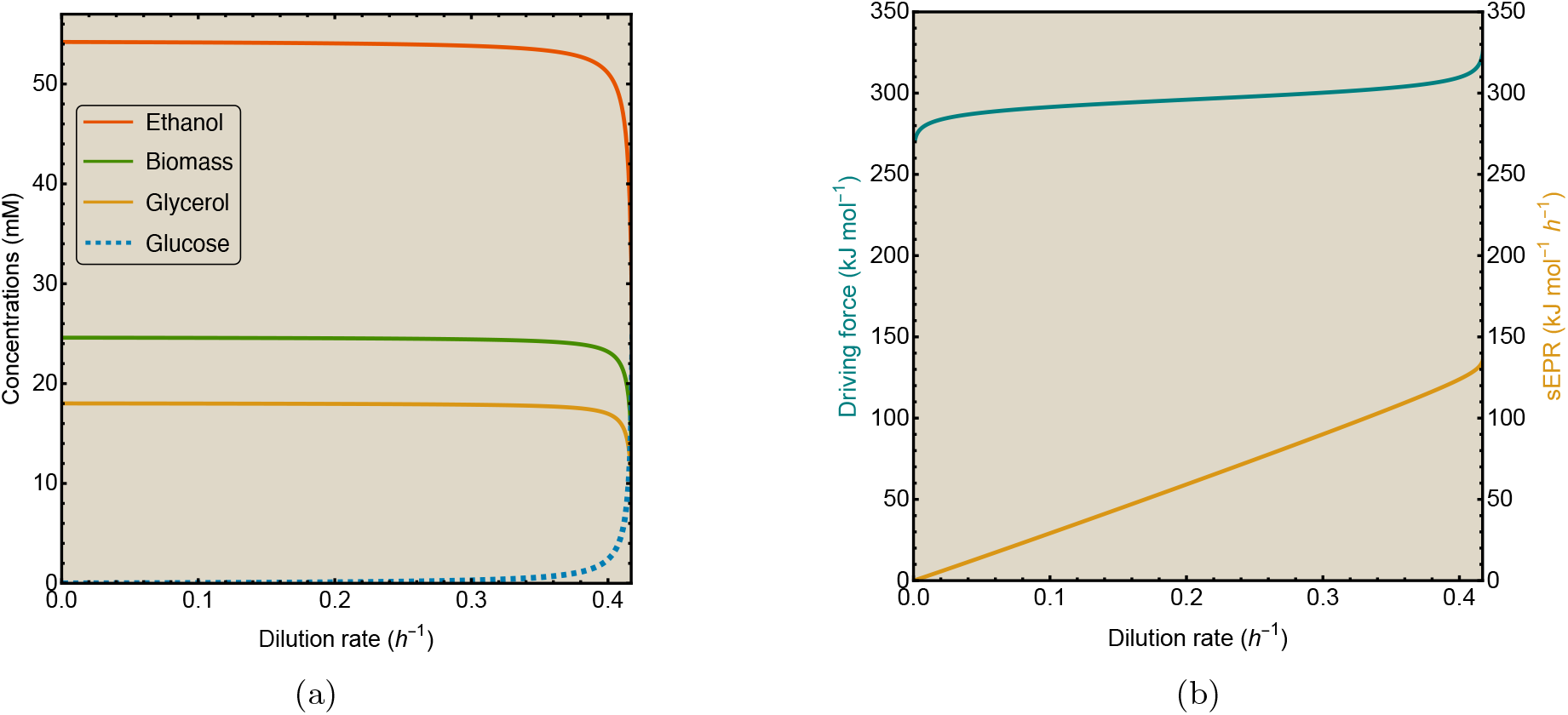
The sEPR rises with growth rate for chemostat growth of anaerobic yeast. (a) Concentrations and (b) Thermodynamic quantities for anaerobic chemostat cultivation of *S. cerevisiae* on glucose. Model parameters are given in Table 1.

When we consider this fixed macrochemical equation in a chemostat model using steady-state equations (17), then we show in Appendix E that the driving force rises with the growth rate. This follows essentially from the way the growth rate (16) depends on substrates and products, and the fact that substrate concentrations increase with the growth rate, while product concentrations decrease. Hence, the sEPR also rises with the growth rate, as is also illustrated by Figure 3b. We also provide a second example, for the Gram-negative bacterium *Klebsiella aerogenes*, with similar results in Appendix D. Parameters for both models are given in Table 1. Mathematica implementation of the chemostat model for different organisms can be found on https://github.com/MaartenJDroste/Entropy-Production-Rate.

**Table 1.** Table with parameter values for each organism for which a chemostat model has been developed. The references next to the organism names are the reference from which its macrochemical equation is taken. The last three cases of *S. cerevisiae* are simulated with a model including overflow metabolism. Concentrations in the reservoir medium are chosen such that only the carbon source, which is the first substrate in the list, is limiting growth. For parameters with the same value for different yeast models that are taken from a reference, only the first mention of the parameter value contains the corresponding reference.

| Organism | Environment | $\lambda_{max}$ (h<sup>-1</sup>) | Reservoir concentrations (mM) | $K_{S_c}$ (mM) | $\Delta\mu_{growth}^o$ (kJ/mol) | $D_c$ (h<sup>-1</sup>) |
| --- | --- | --- | --- | --- | --- | --- |
| *K. aerogenes*<br>[Heijnen and Dijken, 1992] | Anaerobic | 0.62<br>[Streekstra et al., 1987] | $[C_6H_5O_7^{3-}]_R = 100$<br>$[NH_4^+]_R = 100$ | 0.24<br>[Johnson et al., 1975] | -185 [Heijnen and Dijken, 1992] | - |
| *S. cerevisiae*<br>[Battley, 2013] | Anaerobic | 0.42 [van Hoek et al., 1998] | $[C_6H_{12}O_6]_R = 41.7$<br>[van Hoek et al., 1998],<br>$[NH_3]_R = 50$,<br>$[H_3PO_4]_R = 50$,<br>$[H_2SO_4]_R = 50$ | 0.12<br>[Snoep et al., 2009] | -357 [Battley, 2013] | - |
| *S. cerevisiae*<br>(Model 1)<br>[Battley, 2013] | Aerobic | 0.42 | $[C_6H_{12}O_6]_R = 41.7$<br>$[O_2]_R = 250$<br>$[NH_3]_R = 50$<br>$[H_3PO_4]_R = 50$<br>$[H_2SO_4]_R = 50$ | 0.12 | $\Delta\mu_{B,res}^o = -1046$,<br>$\Delta\mu_{B,fer}^o = -357$ | 0.28 |
| *S. cerevisiae*<br>(Model 2)<br>[Heijnen and Dijken, 1992] | Aerobic | 0.42 | $[C_6H_{12}O_6]_R = 41.7$<br>$[O_2]_R = 250$<br>$[NH_4^+]_R = 50$ | 0.12 | $\Delta\mu_{res}^o = -467$,<br>$\Delta\mu_{fer}^o = -255$ | 0.28 |
| *S. cerevisiae*<br>(Model 3) | Aerobic | 0.42 | $[C_6H_{12}O_6]_R = 41.7$<br>$[O_2]_R = 250$<br>$[NH_4^+]_R = 50$<br>$[H]_R = 50$ | 0.12 | $\Delta\mu_{res,1}^o = -330$<br>$\Delta\mu_{ethfer}^o = -294$<br>$\Delta\mu_{acefer,1}^o = -357$<br>$\Delta\mu_{res,2}^o = -326$<br>$\Delta\mu_{acefer,2}^o = -329$ | $D_{c1} = 0.28$<br>$D_{c2} = 0.35$ |

### 3.5. Shifts in metabolic strategies can lead to a reduced sEPR as function of the growth rate in the chemostat

Many microorganisms gradually shift metabolic strategies as function of growth rate in aerobic, nutrient-limited chemostats after a critical dilution rate (*D*_*c*_). For instance, *S. cerevisiae* shifts from glucose respiration to fermentation into ethanol [Elsemman et al., 2022] or *L. lactis* shifts from mixed-acid fermentation (formation of acetate, formate and ethanol) to homolactic fermentation (lactate formation) [Goel et al., 2015]. This implies that the net macrochemical equation of growth is the combination of two such equations (in the absence of maintenance requirements, which are not taken into account in this paper), one for each metabolic strategy, as explained in Section 2.5. The substrate and product yields on biomass are a convex combination of the corresponding quantities of both strategies. Focusing on the shift in *S. cerevisiae*, the yield of metabolite *i* on biomass depends on the dilution rate *D* as

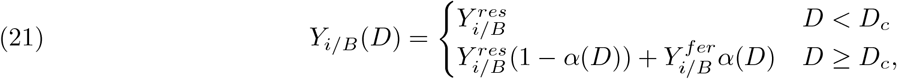

where 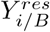 and 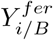 are the yields of metabolite *i* for respiration and fermentation, respectively.

Similarly, the biological^2^ standard Gibbs energy 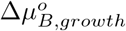 depends on *D* as

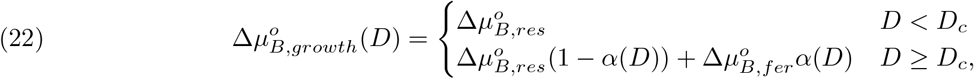

because it is directly determined by the yields. Here 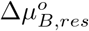 and 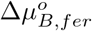 are the biological standard Gibbs energy dissipation of respectively respiration and fermentation.

To describe the Gibbs energy potential Δ*μ*_*growth*_(*D*) including contributions from external concentrations in a similar way, eq. (15) and (22) can be combined. Splitting the logarithmic concentration term in separate contributions for respiration and fermentation results in

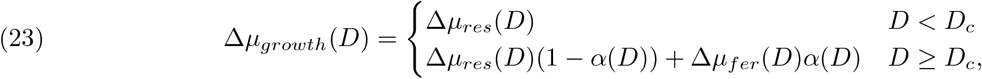

where Δ*μ*_*res*_(*D*) and Δ*μ*_*fer*_(*D*) denote the Gibbs energy potential of respiration and fermentation including contributions from external concentrations, which therefore change with *D*.

In these expressions we introduce a mixing function *α*(*D*), which can be interpreted as the fraction of biosynthetic resources directed to fermentation. So, this mixing function is equivalent to a convex coefficient as in (5), and is therefore also denoted by *α*. Experimental data [Basan et al., 2015, van Hoek et al., 1998] suggests that relations for uptake and excretion rates *q*_*i/B*_(*D*) = *DY*_*i/B*_(*D*) are affine. In Appendix F we show that this implies a mixing function of the form

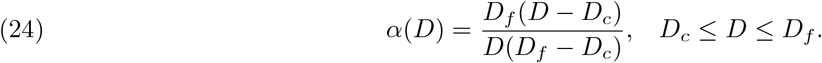

The parameter *D*_*f*_ represents the dilution rate at which the organism would have replaced respiration completely by fermentation, and can therefore be called the ‘final’ dilution rate. It should be noted that *D*_*f*_ is only a fitting parameter in our models and is not an observable dilution rate (it generally exceeds the maximal growth rate in the chemostat, because the microbial culture will have washed out before this final dilution rate is reached). The mixing function (24) attains values between 0 and 1 for *D*_*c*_ ≤ *D* ≤ *D*_*f*_. More information on modeling shifts in metabolism, other possible forms for the mixing function and an extension to mixing three strategies can be found in Appendices F-H.

As the mixing function *α*(*D*) describes mixing of metabolic strategies that changes with dilution rate, we have extended the chemostat model from Section 2.6 to microorganisms with a metabolic shift. We have constructed three different models of the yeast *S. cerevisiae* shifting from glucose respiration to fermentation into ethanol. Each of these models uses the chemostat model as basis, but has different macrochemical equations for respiration and fermentation as main ingredients. For model 1, macrochemical equations from [Battley, 2013] are used as inputs. Battley describes respiratory growth on glucose by *S. cerevisiae* with

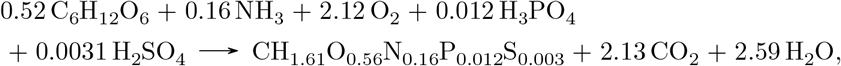

while (20) refers to the macrochemical equation associated with ethanol fermentation. Model 2 uses macrochemical equations for respiration and fermentation of yeast from [Heijnen and Dijken, 1992]. Model 3 uses macrochemical equations that we have constructed by fitting a genome-scale model to experimental chemostat data of [van Hoek et al., 1998]. Model 3 contains five different macrochemical equations, as it has two critical growth rates *D*_*c*1_, *D*_*c*2_ and also acetate excretion by another metabolic strategy. A detailed description of each model including parameters is given in Appendix G and H.

From these input macrochemical equations and the specific flux data of ethanol, as measured by [van Hoek et al., 1998], we can determine the fraction of glucose directed to fermentative growth for each yeast model. The appropriate mixing function for each model is obtained by fitting two different functional forms to these investment fractions. The form (24) based on linear flux dependencies is used first, unless the fitting parameters *D*_*c*_, *D*_*f*_ have unrealistic values. In that case we use a different form for the mixing function based on hyperbolic flux dependencies, which is described in detail in Appendix F.

As an example, Figure 4 shows the metabolite yields and the specific fluxes that are calculated for model 3. Its mixing function(s) are obtained from fitting the appropriate convex combination of the corresponding macrochemical equations to the ethanol and acetate fluxes as measured by [van Hoek et al., 1998]. This should result in specific fluxes for model 3 that are comparable with this experimental data. Indeed, the model fits the specific flux data relatively well, except for oxygen.

**Figure 4.**
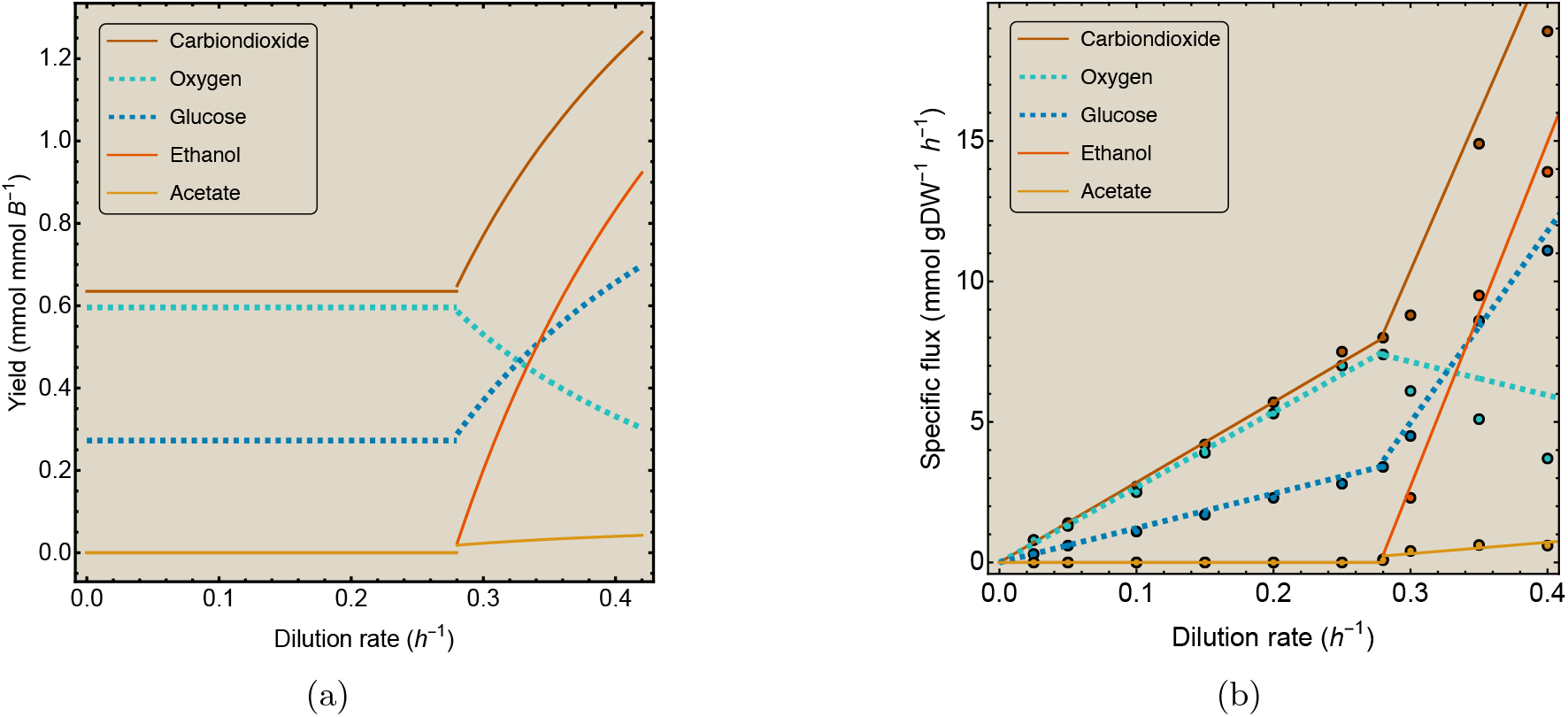
Comparison of model 3 with the data from [van Hoek et al., 1998]. (a) Yields and (b) uptake and excretion rates for model 3 of overflowing *S. cerevisiae*. To determine the mixing of strategies for this model, it is fitted to experimental data for acetate and ethanol flux from [van Hoek et al., 1998], which are represented by dots. The resulting model fluxes are represented as lines. Substrates are shown by dashed lines, products by solid lines.

Equations (21)–(23) are used as ingredients for each yeast model, instead of one macrochemical equation as for the case of a constant metabolic strategy. Together with the steady-state concentrations that are obtained with the chemostat model (12)–(16), these are all ingredients to determine the sEPR for each model, as a function of the dilution rate. The results are shown in Figure 5. Concentrations of only the key chemical compounds are depicted, which for all models show similar qualitative behaviour. They are approximately constant for *D < D*_*c*_ (when the yields remain fixed), but change above the critical dilution rates, when the yields become dependent on the dilution rate. Close to *D*_*max*_, biomass and other products wash out of the vessel, which explains their concentrations approach zero and the glucose concentration rises to the concentration in the medium that flows into the chemostat.

**Figure 5.**
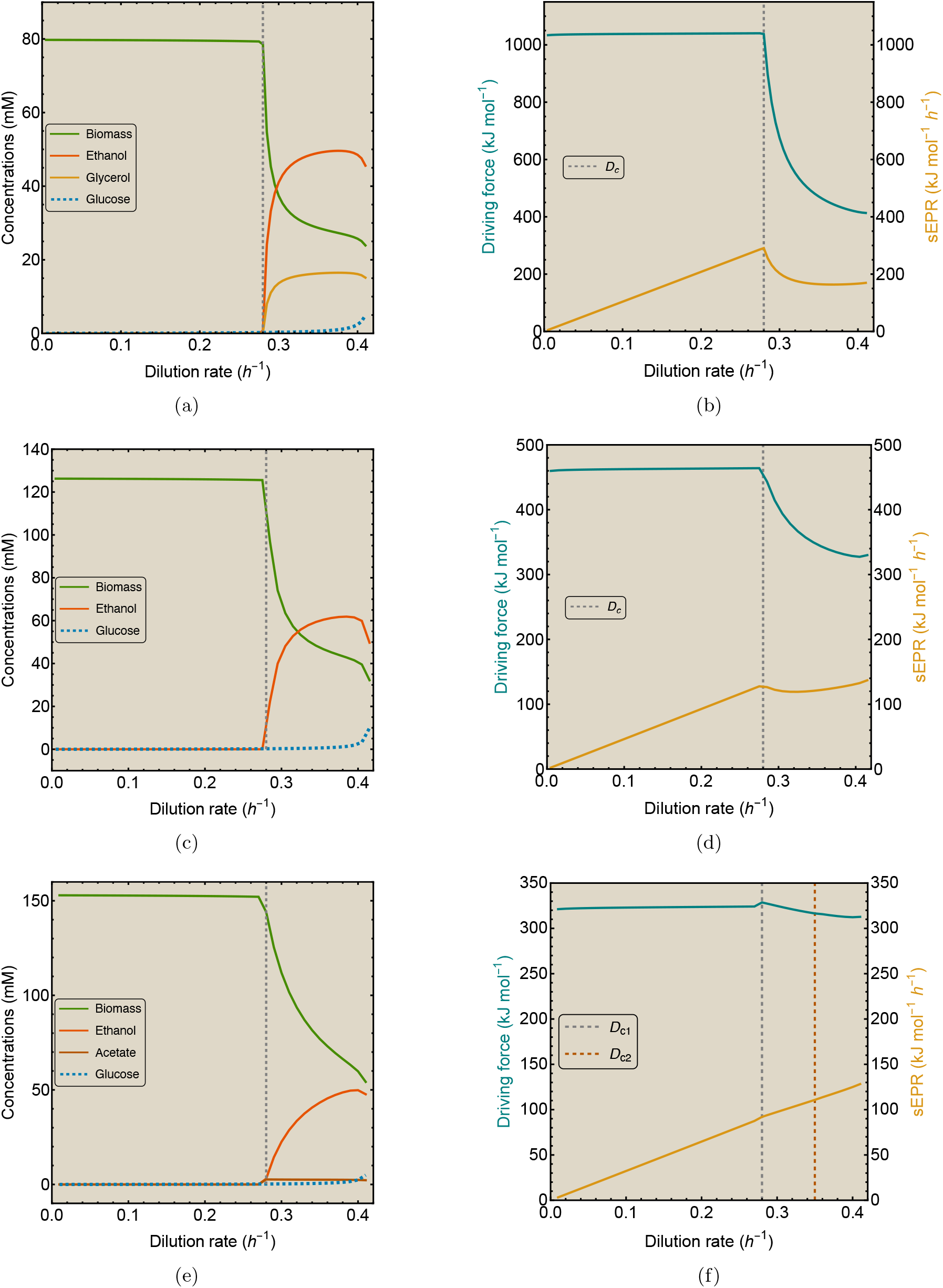
Three chemostat models for overflowing yeast displaying different behaviours of the sEPR. (a) Concentrations and (b) thermodynamic quantities for model 1 for *S. cerevisiae* growing aerobically on glucose, shifting from respiration to fermentation. (c), (d) Same quantities for model 2. (e), (f) Same quantities for model 3. Substrates are shown by dashed lines, products by solid lines. Parameters for each model are given in Table 1.

As respiration degrades glucose further than fermentation, the respiratory driving force is generally larger than that of fermentation; *X*_*res*_ *> X*_*fer*_. Before cells shift from respiration to fermentation, the driving force remains essentially constant for all models, but decreases after the shift, when *D > D*_*c*_. Figure 5 shows that its decrease basically depends on the ratio of standard energies 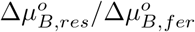 and the mixing function *α*(*D*) for the corresponding model. This results in different b ehaviours of the s EPR for each model.

The sEPR increases monotonically for *D < D*_*c*_, as each yeast model uses only respiration and thus has a fixed macrochemical equation, consistent with Section 3.4. For *D > D* _*c*_, the s EPR of model 1 in Figure 5b decreases as function of the growth rate, while the sEPR of model 2 in Figure 5d only shows a small dip before increasing again. For model 3 in Figure 5f the behaviour of the sEPR hardly changes after critical growth. The second shift in this model does not give any observable effect in the calculated quantities.

These results suggest that as growth rate increases, the shift from respiration to fermentation indeed does not need to coincide with an increasing sEPR. In the next section we derive a quantitative rule of thumb that can be used to predict whether the sEPR decreases after the critical growth rate or not.

### 3.6. A quantitative criterion for prediction of behaviour of the sEPR

The previous results suggest that if the driving force decreases faster than linear with the dilution rate *D* after the shift, the sEPR can decrease when *D* ≥ *D*_*c*_. This can happen either when 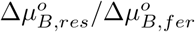 is large, or by a steeply increasing mixing function *α*(*D*) with *D*.

When growth is far from thermodynamic equilibrium, the concentration effect on the Gibbs energy change can generally be neglected, i.e., 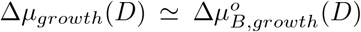. If the mixing function *α*(*D*) is of the form (24), we can then approximate the sEPR by 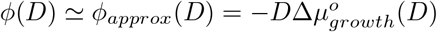 (see Appendix I).

During a shift from metabolic strategy 1 to strategy 2 for *D > D*_*c*_, a decreasing *ϕ*_*approx*_(*D*) with *D* occurs now when the following criterion is met:

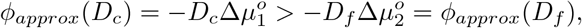

i.e., when,

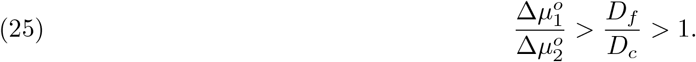

This criterion indicates that a microbe should shift to a strategy with a much smaller driving force, or this shift should occur in a small range of dilution rates to get a decreasing sEPR.

The criterion depends on only a few model parameters and is independent of the dilution rate. It can be evaluated for each chemostat model of a microbe displaying a metabolic shift, without explicitly running the complete chemostat model and knowing all other model parameters. The four parameters in (25) can be plotted as 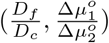. If this coordinate lies above the diagonal given by 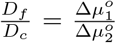, then the sEPR is predicted to decrease with growth rate. Figure 6 shows these parameter combinations for the three models discussed in the previous section as blue dots. For model 1 and 2, the sEPR is predicted to decrease, while for model 3 an increase is predicted; these predictions are in line with Figure 5 for model 1 and 3 but not for model 2. Appendix J provides a detailed description of the criterion prediction for each yeast model.

**Figure 6.**
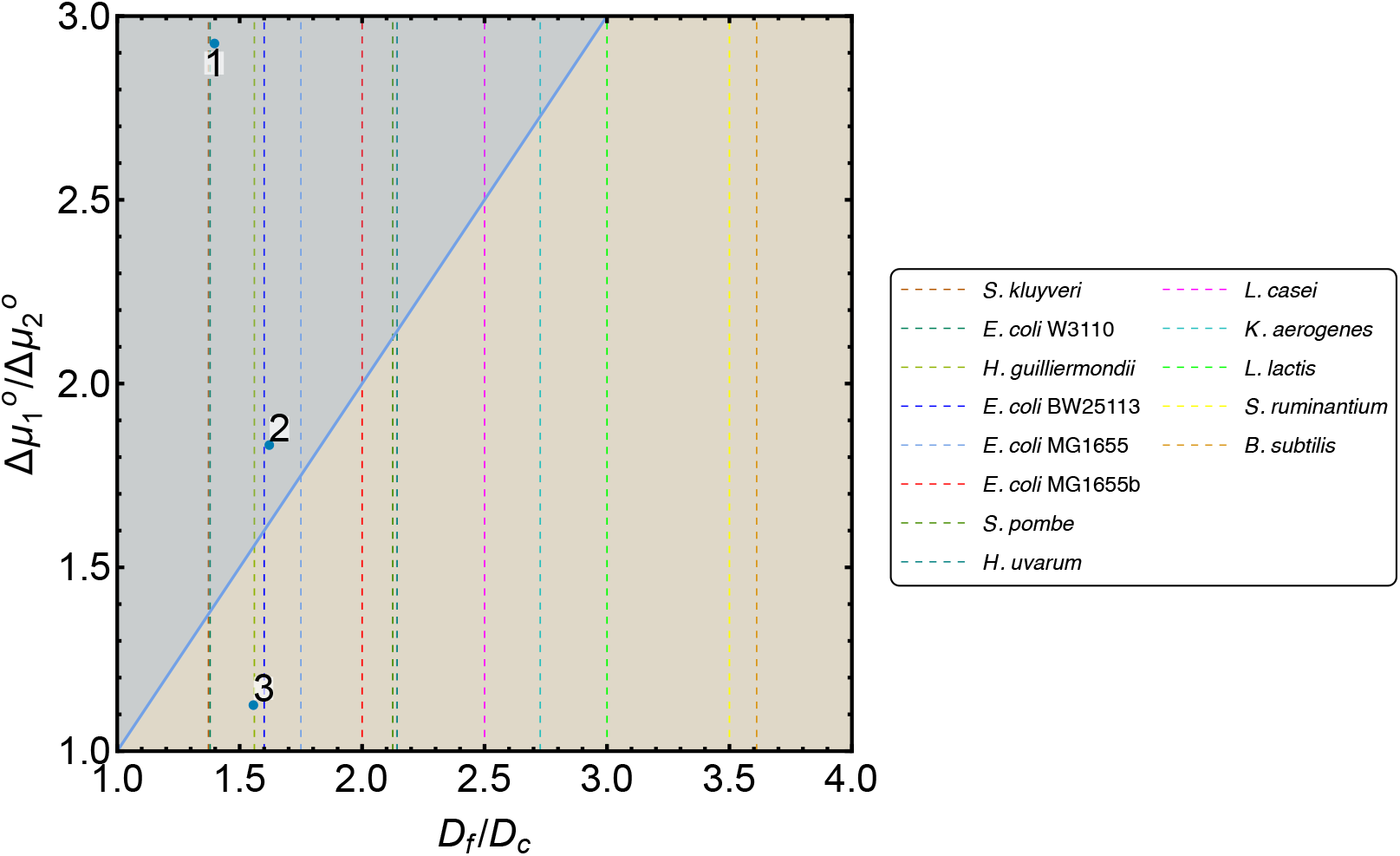
A quantitative criterion can predict the behaviour of the sEPR. The sEPR decreases in the blue region. Blue dots represent the three models from this work. Vertical lines depict growth data for metabolic shifts in different other microbes. The growth data is given in Table 2.

We have tried to apply the criterion to other organisms that display a metabolic shift. Critical and maximal/final dilution rates have been reported for several species, but we generally lack experimental data to characterise the ratio of driving forces between the two metabolic strategies. We have plotted these cases using vertical dashed lines in Figure 6.

**Table 2.**
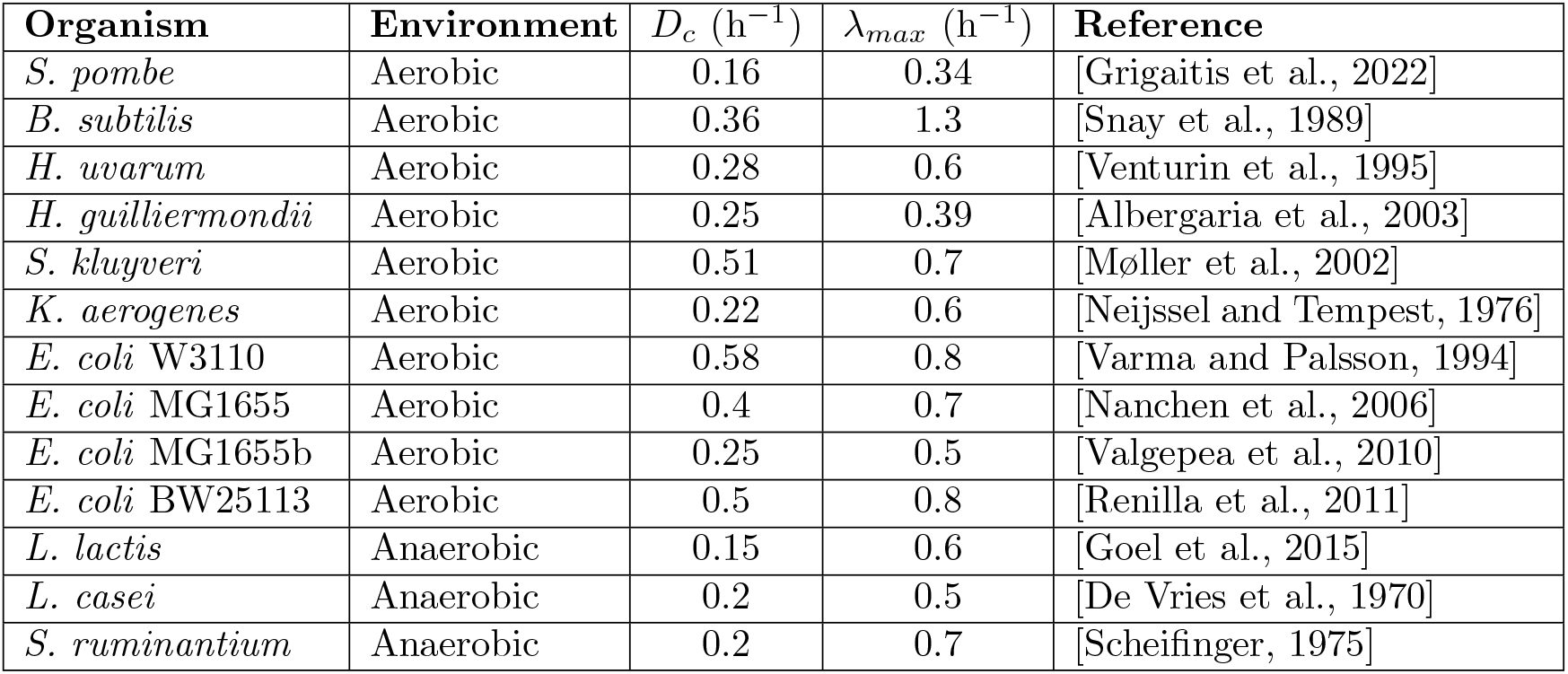
Table with growth data for organisms displaying a shift in metabolic strategies.

The organisms *L. lactis, L. casei* and *S. ruminantium* show metabolic shifts during anaerobic growth. For these and other anaerobic shifts, catabolic Gibbs energy potentials are well-characterised for both strategies, but total Gibbs energies of both strategies can not be determined as the coupling to anabolism and its Gibbs energy potential can only be approximated crudely. The other depicted organisms show metabolic shifts in aerobic environments, which are similar to overflow metabolism in yeast. For these organisms, data on critical growth rates *D*_*c*_ is available, and *λ*_*max*_ can be used as a lower bound for *D*_*f*_. Data for the standard free energy potential 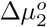 of the second strategy used by the organism is not available.

All available data is presented in Table 2. Details about the exact metabolic shift, the corresponding growth data and other information for each organism are described in Appendix J.

## 4. Discussion & Conclusion

In this work we have studied the relation between entropy production rate (EPR) and growth rate in microbial cells. We have demonstrated that the EPR of a chemical reaction network with a fixed macrochemical equation rises with its reaction rates, because of the direct relation between rate and thermodynamic driving force. Metabolic networks in living cells have more degrees of freedom: they can change enzyme concentrations, allowing them to choose different reactions and alter reaction rates. This may result in decreasing EPR at faster growth, due to a shift towards a metabolic strategy with a lower driving force.

Studying balanced microbial growth in chemostats allows for controlled analysis of both the growth rate and corresponding (s)EPR. By extending a chemostat model with thermodynamics we have gained insight in the behaviour of the EPR as function of growth rate. We have given sufficient conditions for the sEPR to rise with growth rate. For steady-state growth of microorganisms in a chemostat, this is true whenever the net conversion of substrates into products—the macrochemical equation—is fixed.

However, some microbes shift from an energetically-efficient to an energetically-inefficient strategy at faster growth. Such a metabolic shift changes the net conversion of the metabolic network, as the inefficient strategy only partially degrades the carbon source (e.g., glucose) to e.g., acetate, ethanol or lactate. This partial degradation extracts less free energy and can therefore have a lower driving force at higher growth rates. In general, microbes can compensate for the resulting rate loss through reinvestment of biosynthetic resources into protein expression. A smaller, energetically-inefficient pathway can function at higher enzyme concentrations than a larger, energetically-efficient one, by investing the same overall protein concentration in fewer reactions. In this way, the reduction in driving force is compensated by higher rates. This leads to a higher growth rate, but the sEPR does not necessarily increase as well. We further illustrate this using numerical calculations of three different chemostat models for the yeast *S. cerevisiae* that exhibits overflow metabolism at high glucose availability in aerobic environments.

The (standard) Gibbs free energy dissipation during aerobic growth on glucose is reported to be 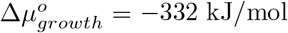 [Heijnen and Dijken, 1992]. This value is consistent with the Gibbs energy dissipation around *D* = 0.4 h^−1^ in both model 2 and model 3, see Figures 5d and 5f. Both these models mix different macrochemical equations, but show very different behaviour in their sEPR, even though they have a similar driving force at high growth rates. Model 3 is however based on a GEM that uses constraint-based methods based on data and contains a large part of the metabolic network of yeast. One could therefore argue that model 3 gives more reliable results.

To make more concrete statements, we need additional experimental data confirming our theoretical predictions. There have been calorimetric measurements of heat production in chemostats [von Stockar and Birou, 1989, Ishikawa and Shoda, 1983], but none of these are suited to compare with the results displayed here. We think this is, next to the difficulty of measuring EPR, because only a few authors consider chemostats as an interesting setup to study thermodynamic properties of living systems. Since experimental evidence is missing and our models show different results, it is too early to state that the sEPR of *S. cerevisiae* will decrease above the critical growth rate. However, the results for all three models show a decreasing driving force as growth rate increases. Only in specific cases when the decrease in driving force compensates for the increase in growth rate during the shift, this will also result in a decreasing sEPR as growth rate rises. This occurs in one of our chemostat models for overflowing yeast. Every situation in which microbes perform a metabolic shift thus has to be assessed separately, at present.

To give a prediction whether sEPR goes up or down as the growth rate passes a critical value at which a metabolic switch occurs, we have given a simple quantitative rule of thumb, see eq. (25). This criterion involves only a small number of measurable parameters: the ratio of critical and final dilution rates, and the ratio of standard Gibbs free energy potentials for the different strategies. However, it turns out that even these parameters have not been experimentally determined for most organisms. To further expose thermodynamic properties of microbes and their metabolic shifts, these parameters should be determined experimentally during chemostat growth. This requires measurements of substrate and product concentrations and fluxes, and calorimetric measurements of heat fluxes. A different method would be to derive macrochemical equations from GEMs fitted to chemostat data, as is done in this work.

For out-of-equilibrium physicochemical systems the EPR is an important quantity, as it describes the free energy dissipation rate and is therefore a measure for the distance from thermodynamic equilibrium [Dilip Kondepudi and Prigogine, 2014, Martyushev and Seleznev, 2006]. Since living systems belong to this class, the EPR has been studied in the context of evolution [Jennings et al., 2020], ecosystems [Meysman and Bruers, 2010] and aging [Balmer, 1982, Yildiz et al., 2020]. It has been suggested by various authors that EPR can be used to understand microbial growth, as it has been interpreted as a driver for microbial evolution [Unrean and Srienc, 2011], or it appears to reach an upper limit that governs cellular metabolism [Niebel et al., 2019].

On the other hand, other studies show that microbial behaviour can be understood from principles of resource allocation to attain high growth rates (reviewed in [Bruggeman et al., 2020]). In this work we have demonstrated that EPR can be interpreted as thermodynamic measure for microbial growth rate in a chemostat, but that this relation can break down if the unicellular organism shifts towards a metabolic strategy with lower driving force at faster growth. We therefore tentatively conclude that EPR can not be used as a reliable predictor of microbial growth.

The thermodynamic chemostat model developed in this work can also be used to study other organisms showing different exotic shifting behaviour in the chemostat. Since it only requires knowledge of macrochemical reactions and a few measurable parameters, it is fairly easy to extend. It is therefore an example of the usefulness of the thermodynamic black-box approach, as also highlighted by others [Westerhoff et al., 1982, Qian and Beard, 2005, von Stockar et al., 2006, Saadat et al., 2020]. However, to understand what determines the behaviour of the (s)EPR during metabolic shifts, we and others [Ebenhöh et al., 2023] believe more detailed models of metabolism are required that decompose growth in a catabolic and anabolic process and include their energy coupling, analoguous to the energy converter principle [Wilken et al., 2021]. This will be investigated in future work. Furthermore, we considered microbes growing far from thermodynamic equilibrium, but there are also cases of growth close to equilibrium [Allaart et al., 2023], which might show different behaviour of the sEPR. Lastly, our modeling approach opens the door for other thermodynamic analyses of microbes growing in different environments, such as batch conditions. We hope that this work stimulates further research in microbial physiology using thermodynamics, in theoretical as well as experimental directions.

## 5. Glossary

*b*: Biomass concentration in the chemostat vessel.
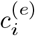: (Equilibrium) Concentration of chemical compound *C*_*i*_. *D*: Dilution rate of a chemostat.
*D*_*c*_: Critical growth/dilution rate after which a shift in metabolic strategies occurs.
*D*_*f*_: Dilution rate at which an organism has replaced its metabolic strategy at slow growth completely by its strategy at fast growth.
*D*_*max*_: Maximal dilution rate in a chemostat, above which wash-out of cells is faster than growth. For 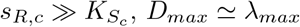.
*e*_*j*_: Concentration of enzyme *j* catalysing reaction *j*.
**E**_*i*_: Elementary flux mode *i* of a metabolic network.
*e*_*T*_: The concentration of all enzymes in a metabolic network summed together.
*f*_*j*_(**c**^*′*^): Saturation function of enzyme *j* catalysing reaction *j*.
j: Vector containing the number of moles of reactants consumed and/or produced in the reactions, obtained from normalising the flux vector 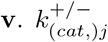Forward/backward (catalytic) rate constants of reaction *j*.
*K*_*eq,j*_: Equilibrium constant of reaction *j*, directly related to its standard Gibbs energy dissipation.
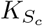: Monod saturation or affinity constant for carbon source *S*_*c*_ in the chemostat.
N: Stoichiometric matrix of a chemical reaction network with entries *n*_*ij*_.
N^*′*^: Stoichiometric matrix extended with stoichiometric coefficients of externally fixed substrate and product concentrations.
*p*_*l*_: Concentration of product *P*_*l*_ in the chemostat vessel.
*q*_*i/B*_: Uptake or excretion rate of compound *C*_*i*_ per mole biomass per hour.
*S*_*c*_: Carbon source, usually limiting growth.
*s*_*k*_: Concentration of substrate *S*_*k*_ in the chemostat vessel. *s*_*R,k*_: Concentration of substrate *S*_*k*_ in the reservoir medium. *R*: The universal gas constant, 8.314 J/(mol K).
*T*: Temperature of the system.
*v*_*j*_: Rate of reaction *j*.
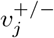: Forward/backward reaction rates.
*X*: The thermodynamic driving force of a reaction or pathway per mole, which equals minus the Gibbs free energy potential.
*Y*_*i/B*_: Yield in moles of metabolite *C*_*i*_ per mole biomass.
*α*_*i*_: Conic coefficients for a flux vector decomposition in terms of the EFMs of the reaction network.
*α*(*D*): Mixing function representing the fraction of resources invested in fermentation.
Δ*μ*: The Gibbs free energy potential per mole of a reaction or pathway.
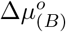: The (biological) standard Gibbs free energy potential per mole of a reaction or pathway.
*λ*: Growth rate of a microbial culture.
*λ*_*max*_: Maximal growth rate (during batch cultivation).
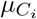: Gibbs free energy per mole of compound *C*_*i*_.
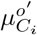: Standard Gibbs free energy per mole of compound *C*_*i*_.
Φ: Entropy production rate (EPR).
*ϕ*: Specific entropy production rate (sEPR) scaled with temperature.
*ϕ*_*Approx*_: Approximation of the sEPR that neglects concentration effects.

## Acknowledgements

We would like to thank our colleagues of the VU Systems Biology Lab for critical and stimulating discussions. Also, we thank Robbert Kleerebezem and Elad Noor for their help in understanding and applying thermodynamic analyses to (models for) different microorganisms. MJD and RP acknowledge funding by NWO grant 613.009.131 ‘Control of maximal growth rate by single-celled organisms’. MR and FJB acknowledge funding by NWO-XL grant OCENW.XL21.XL21.007 ‘Taking Control of Metabolism in Microbial Cell Factories by Applying Noncanonical Redox Cofactors’.

## Appendix A. Considerations for Gibbs energies

For any reaction *j*, the molar Gibbs free energy potential can be expressed as

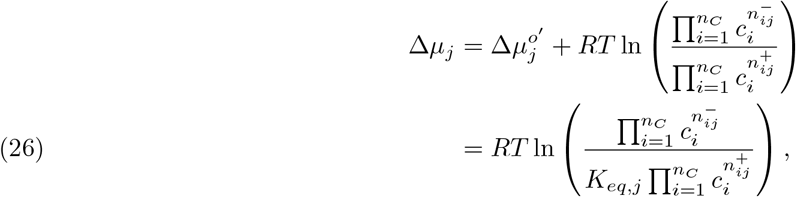

where the equilibrium constant of the reaction is

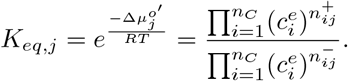

This relation implies that the reaction is in thermodynamic equilibrium Δ*μ*_*j*_ = 0 when all concentrations are equal to their equilibrium concentrations 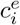, making the Gibbs energy dissipation a measure for the distance from thermodynamic equilibrium. The standard Gibbs energy of reaction 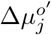 can therefore be derived directly from this equilibrium condition, as it represents the change in Gibbs free energy occurring at standard conditions.

Standard conditions represent a reference state, denoted by the superscript ^*o*^, which has all concentrations at 1 mol/L, a pH of 7, pressure of 1 bar and a temperature of 298.15 K. For biological systems, concentrations of 1 mmol/L are more realistic. These biological standard conditions, which are mostly used in this work, are therefore indicated by an extra subscript _*B*_. Note that this changes the reference state and thereby also the standard Gibbs energies (and the corresponding equilibrium constants), but not the values of the Gibbs energies as calculated by eq. (26). Furthermore, as microbes live in an aqueous environment, all reactions occur in an aqueous solution which changes the corresponding Gibbs energies as well. This is usually denoted by a prime ‘, but this notation is suppressed in the rest of this work as we only consider aqueous environments. Moreover, the water concentration hardly changes during microbial growth. Therefore, water is not taken into account in calculating Gibbs energy changes, as it remains at its equilibrium concentration.

More information on these different standard conditions and their implications can be found on the eQuilibrator website [Flamholz et al., 2011]. This is a useful tool for calculating standard Gibbs energies in different conditions. A complete analysis of the effects of a change in standard conditions on the corresponding Δ*μ*^*o*^ is performed by Popovic [Popovic, 2019].

To calculate 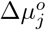 for a chemical reaction, the formation energies 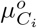 at standard conditions of all reactants need to be known, which can be calculated with methods like component contributions [Noor et al., 2013].

They are tabulated in different studies of thermodynamics of microbial growth [Heijnen and Dijken, 1992, Battley, 2013, Alberty, 1998]. These formation energies represent the change in Gibbs free energy during the formation reaction of one mole of the compound. Just as the Gibbs energy dissipation Δ*μ*_*j*_ can be expressed in terms of molar Gibbs free energies of the reactants, 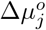 can be calculated as

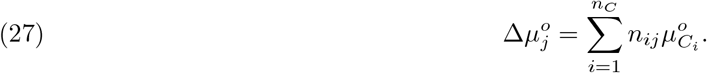

It is common practice to define a reference frame for the formation energies by setting some of them to zero, which can be convenient during calculations or measurements. However, this makes it harder to compare literature values for formation energies, as these are different in each reference frame. This also changes the corresponding Gibbs energies of reaction, which makes it important to be consistent in the use of formation energy tables during modeling.

In the chemostat models constructed in this work, 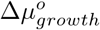 is a parameter for each macrochemical equation. For macrochemical equations extracted from the literature the value for 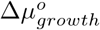 is given in the corresponding reference. All values are such that growth is always far-from-equilibrium. Recalculation shows that indeed most references use different standard conditions and reference frames, to which we changed our simulations consistently. Details for our models are described in Appendices D, G and H.

Because of the difference in standard conditions and reference frames, we performed a check by calculating 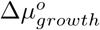 for each model with the table from Heijnen and Kleerebezem [2010], which uses the thermodynamic reference frame. This is the most complete table we could find and therefore serves well as a calculation check. The resulting 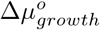 for the macrochemical equations from Heijnen and Dijken [1992] in model 2 only have a maximum difference of 3%, as their table is almost completely comparable to the table in [Heijnen and Kleerebezem, 2010]. For the macrochemical equations from Battley [2013] in model 1 the maximum difference in 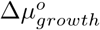 was about 19%. This large discrepancy can however be attributed to the difference in standard conditions. For example, Battley uses standard concentrations of 1 mM instead of 1 M. Therefore a fair comparison of 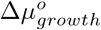 values calculated with these different tables is actually not possible. Also, it should be noted that these differences in 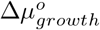 do not affect the qualitative behaviour of the sEPR in Figure 5.

## Appendix B. Mathematical proof for increase of flux and EPR as function of the driving force in chemical reaction networks

Here we present a mathematically precise version of the argument made in Section 3.1. This argument states that the EPR and flux always increase with thermodynamic driving force in a chemical reaction network with mass action kinetics rate laws, provided that the net conversion of substrates into products remains fixed.

Consider a chemical reaction network at fixed pressure and temperature in a non-equilibrium steady-state. It consists of substrates **S**, internal metabolites **C** and products **P** with respective steady-state concentrations **s, c, p**. The conversion

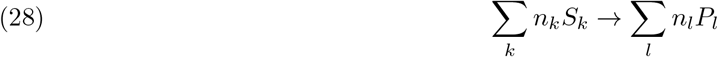

is fixed. The matrix **N**^*′*^ encodes the of stoichiometry the network, while **N** is the submatrix of **N**^*′*^ that contains the rows for the internal metabolites only. The network contains *r* reactions, each reaction *j* running at rate *v*_*j*_(***σ***_*j*_, ***ρ***_*j*_) depending on the substrate concentrations ***σ***_*j*_ and product concentrations ***ρ***_*j*_ of the reaction.

We assume that the reactions occur in a well-mixed medium, so that the rates *v*_*j*_ are determined by the law of mass-action and can be written as

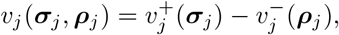

with

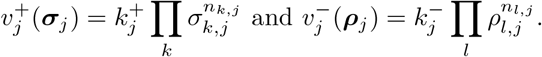

Now we claim that increasing one or more substrate concentrations *s*_*k*_ or decreasing one or more product concentrations *p*_*l*_ results in both a higher flux *v*_*r*_ through the network, and also a higher EPR.

*Proof*. As discussed in the main text, the Gibbs energy dissipation corresponding to reaction *j* is

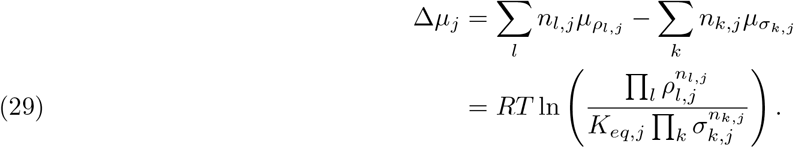

If the network forms an EFM, it has a flux vector **E** that may be normalised such that

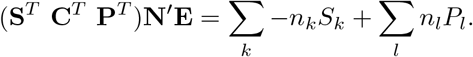

The total Gibbs energy dissipation corresponding to the conversion (28) is then given by

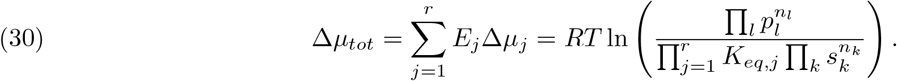

The corresponding driving forces are now given by *X*_*j*_ = −Δ*μ*_*j*_, and *X*_*tot*_ = −Δ*μ*_*tot*_.

As the products of the net conversion are fixed, we may assume that any steady state flux distribution contains a nonzero flux through reaction *r* (producing one of those products). The EFMs of the network {**E**_*i*_}_*i*_ are normalised at their last entries *r*, i.e., *E*_*i,r*_ = 1 for all *i*. For a steady-state flux vector of this network satisfying **Nv** = **0** we may thus also introduce **w** as the corresponding vector with last element 1, i.e., satisfying

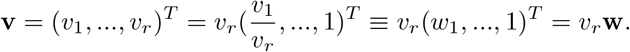

This vector **w** can be written as a convex combination 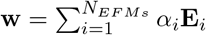 of EFMs with 0 ≤ *α*_i_ ≤ 1 for all *i* and 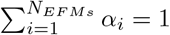 [Gagneur and Klamt, 2004]. This decomposition is non-unique [Schwartz and Kanehisa, 2006]. For the steady-state flux vector **v** this yields

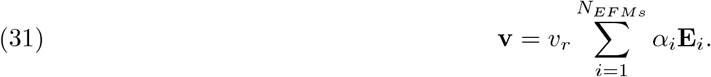

By (30), each EFM has driving force 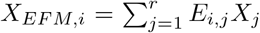. This implies that the driving force of **v** can be written as

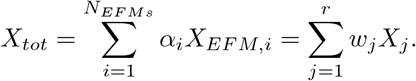

So the driving force is given by a similar convex sum as the flux vector. The EPR of the chemical reaction network is now given by

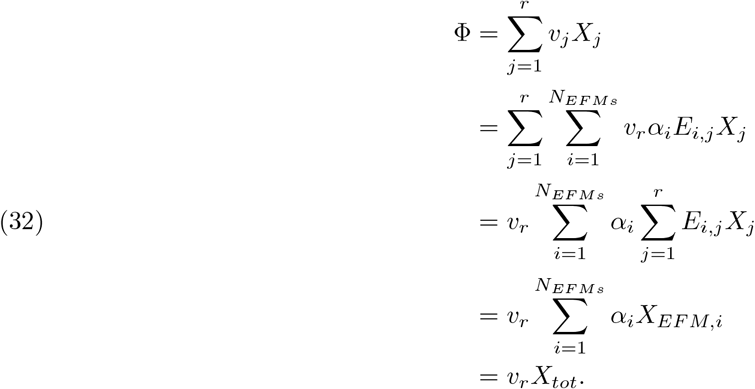

Suppose now that we increase a substrate concentration *s*_*k*_ or decrease a product concentration (or both). Then by (30) the driving force of each EFM *X*_*EF M,i*_ that converts substrates into products according to (28) increases, and hence so does *X*_*tot*_. Moreover, since 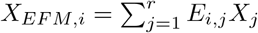, the driving force of at least one reaction *j* must be higher than before. With the assumed mass action kinetics, this reaction thus has a higher flux than before. But in this EFM, all fluxes have fixed ratios, so all fluxes must have increased. We conclude that *v*_*r*_ has increased, and by (32) also the EPR. □

It should be noted that this proof assumes that all EFMs of the reaction network have the same net conversion (28). This condition can be relaxed, if we assume that the driving forces *X*_*EF M,i*_ of all EFMs increase, irrespective of the net conversion by that EFM. This could be realized by increasing all substrate concentrations or decreasing all product concentrations. Assuming^3^ furthermore that every EFM contains reaction *r* and thus has nonzero *v*_*r*_ results in a rising EPR. This follows from a similar argument as in the proof above, as all fluxes and all driving forces increase, so both *v*_*r*_ and *X*_*tot*_ have increased as well.

This version of the proof allows for more freedom in the flux vector, as different EFMs can have different conversions. The proof above allows for more freedom in the external concentrations, as only one substrate or product concentrations needs to change to obtain an increasing driving force *X*_*tot*_.

## Appendix C. Thermodynamic consistent construction of a toy metabolic network

### C.1. General considerations for constructing a toy metabolic network

Small example networks are useful to describe and understand essential properties of metabolic networks, without having to worry about intricate biochemical details. Although this makes the analysis much easier, designing a toy network can be error prone. Kinetic parameters are often chosen freely, while in fact they are restricted by biochemical constraints and the laws of thermodynamics. So, a toy network should be constructed while keeping these rules in mind, a procedure that is sometimes called thermodynamic parameter balancing [Lubitz et al., 2010]. Here we summarise the most important considerations for constructing toy metabolic networks.

First of all, thr network should satisfy the basic relations as described in Section 2. Internal metabolites are in steady-state, represented by ċ = **Nv** = **0**. The rate *v*_*j*_ at which an enzyme-catalysed reaction *j* runs from *S* to *P* in the network is given by some enzyme-kinetic relation 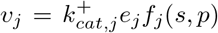. For reversible

Briggs-Haldane kinetics, which is commonly used, the saturation function is given by

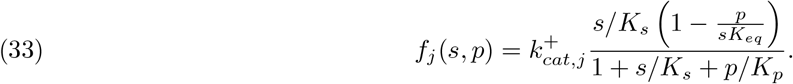

It is also possible to decompose the rate *v*_*j*_ in a forward and backward rate as in eq (2). However, the convenient property of notation (33) is that it automatically satisfies the Haldane relation [Haldane, 1930], which for this type of kinetics can be written as

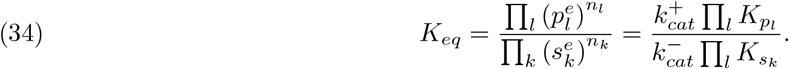

This relation implies a constraint on the kinetic parameters and affinities of every reaction, reducing thereby the degrees of freedom. Eq. (33) is written such that the backward catalytic rate constant 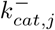 is not included, leaving this as free parameter to tune in the model such that the Haldane relation is satisfied.

If the network contains a closed cycle, this has extra implications. As a closed cycle does not result in any net conversion (there are no external concentrations), its Gibbs energy dissipation (i.e., the total driving force) should vanish: Δ*μ*_*cycle*_ = Σ_*j*_ Δ*μ*_*j*_ = 0. It can be shown that this condition is equivalent to Π_*j*_ *K*_*eq,j*_ = 1. This constraint on Gibbs energies or equilibrium constants is called Kirchhoff’s l oop law, analogous to his law for electrical circuits [Qian et al., 2003]. This condition holds when the network is in steady-state, hence also when it is in thermodynamic equilibrium. In the later case this condition is called detailed balance.

### C.2 Specific considerations for our branched pathway e xample

The general constraints above can be translated to our example network in Figure 1. The values of the parameters used in the numerical implementation of this network to create Figure 2 satisfy all constraints. A Mathematica implementation of the network can be found on https://github.com/MaartenJDroste/Entropy-Production-Rate.

The network has two EFMs with fluxes *J*_1_, *J* _2_ and corresponding driving forces 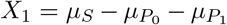 and 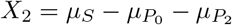. As these depend only on external concentrations, a relation between standard molar free energies of the external metabolites and equilibrium constants of the reactions can be derived for each pathway, using eq. (27). These are given by

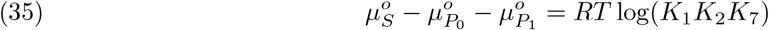

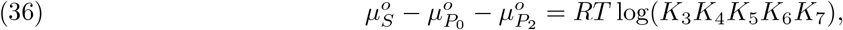

with *K*_*j*_ = *K*_*eq,j*_ for notational convenience. Subtracting the first from the second equation gives

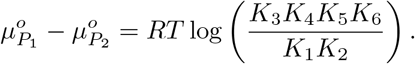

This relation is of course automatically satisfied whenever the others are, but it does show another possible conversion in this example network. Indeed, due to reversibility of all reactions, *P*_2_ can be converted into *P*_1_, and vice versa. This pathway contains an internal cycle, but Kirchhoff’s loop law does not apply here as the cycle is not closed: there are still external metabolites *P*_1_ and *P*_2_, whose concentrations are fixed [Qian et al., 2003].

Comparing the driving forces of EFM 1 and 2 yields

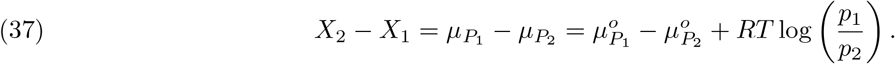

Mass conservation implies that *P*_1_ and *P*_2_ have the same elemental composition, so they are isomers. If *P*_1_ and *P*_2_ are in fact the same molecule, then 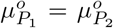. In this case, both pathways perform the same net conversion, so also the concentrations of *P*_1_ and *P*_2_ should be equal, resulting in the same driving force for both pathways as concentrations of *S* and *P*_0_ were already fixed. Also, the network then contains a closed cycle *P*_2_ ⇌ *P*_1_, which satisfies Kirchhoff’s loop law as its driving force vanishes, or equivalently 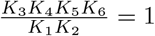. When constructing a toy network these relations are constraints that need to be satisfied by choosing appropriate parameter values.

When constructing a toy network one can prescribe either concentrations and equilibrium constants, or molar free energies and standard Gibbs energies of reactions to satisfy the constraints. As can be seen from the direct relations between these variables the concentration and thermodynamic descriptions are equivalent. Hence, a model in terms of one set of variables and corresponding parameters suffices. This emphasizes that thermodynamics does not encompass any new information regarding modeling.

The branched pathway considered here serves as an example that as function of the nutrient concentration the cell can shift to a pathway that has a higher flux but a lower driving force. To obtain this result, parameters of both pathways can be tuned while satisfying the aforementioned relations and constraints. As EFM 1 has fewer reactions, it has higher (optimal) enzyme concentrations per step so in general also a higher optimal flux when all catalytic rate constants are comparable. Setting the catalytic rate constants of EFM 1 smaller than the rate constants of EFM 2 brings the optimal fluxes through both pathways closer. As explained earlier, this does not violate the Haldane relations because of the remaining degrees of freedom.

Eq. (37) specifies which parameters determine the driving force difference *X*_2_ −*X*_1_. This difference should be positive, which occurs when 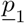 is high while 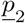 is low (compared to their equilibrium concentrations).

Another way to obtain this is when 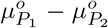 is large, which happens when the equilibrium constants of the reactions in EFM 1 are smaller than the equilibrium constants of the reactions in EFM 2. This can be interpreted as pathway EFM 1 operating closer to thermodynamic equilibrium, and thereby experiencing a larger product inhibition effect. So, the general procedure is to set *K*_*eq,j*_ and *k*_*cat,j*_ for EFM 1 to low values compared to *K*_*eq,j*_ and *k*_*cat,j*_ for EFM 2.

It is even possible to find parameters such that *J*_1_ *> J*_2_, but *X*_1_*J*_1_ *< X*_2_*J*_2_. This case is given by the condition

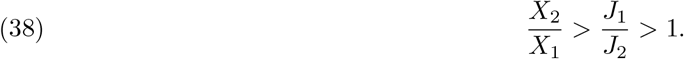

Note that this condition is equivalent to the criterion presented in eq. (25).

## Appendix D. Notes on chemostat modeling & models for two anaerobic microbes

Here we give a detailed description of the models of two cultures of microorganisms growing anaerobically on glucose. The chemostat model equations are given in Section 2.6. A chemostat volume of *V* = 1 liter is used, such that the flow rate *F* equals the dilution rate *D* = *F/V* = *F*. For each organism, its macrochemical equation is the main model ingredient. Here we detail the choices of these equations for the different scenarios discussed in the text.

According to [Battley, 2013], yeast *Saccharomyces cerevisiae* ferments glucose to ethanol and glycerol as

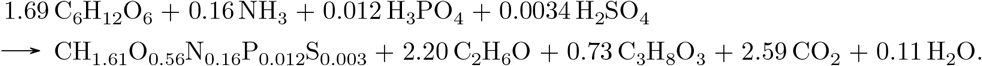

The bacterium *Klebsiella aerogenes* grows anaerobically on citrate, fermenting this to acetate, succinate and formate via [Heijnen and Dijken, 1992]

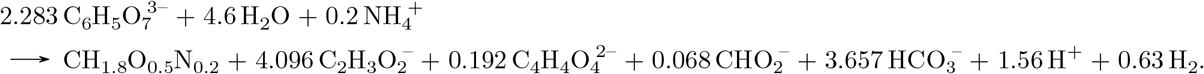

The first product in the macrochemical equation is the biomass, written down in its elemental composition and normalised per C-mole. Most experimental data report the biomass per gram dry weight, which changes the yield units accordingly. Here we adopt the convention of reporting the biomass in moles, to check elemental balance more easily. All model parameters differ per organism and are given in Table 1. The standard Gibbs energies of the macrochemical equations 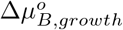 are calculated using the same standard conditions, reference point and thermodynamic tables as in the work stating the particular macrochemical equation. Note that all substrate concentrations in the reservoir medium are chosen such that only the carbon source is limiting for growth. All other substrates are therefore in excess. Also, the concentration of the carbon source in the reservoir medium *s*_*R,c*_ is chosen such that it is much larger than its affinity constant 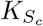.

The macrochemical equations and parameters given in Table 1 are all the ingredients required to implement and numerically solve the chemostat model for these organisms in Mathematica. The numerical solution is calculated by rewriting eq. (17) for the steady-state concentrations and the dilution rate in terms of biomass *b*, and solving this equation for *b* for 0 ≤ *D* ≤ *λ*_*max*_ with a stepsize of 0.001-0.015 h^−1^ (differs per model).

The values *b*(*D*) are then substituted in the other equations to find all other concentrations and thereby also the driving force and the sEPR.

The concentrations and thermodynamic quantities for *Klebsiella aerogenes* are depicted in Figure 7 and are qualitatively similar to the results for yeast in Figure 3. Again, only concentrations of carbohydrates are shown for visual clarity, but other concentrations are included in the models as well and depend similarly on *D*. Concentrations and therefore driving force change only very close to the maximal dilution rate *D*_*max*_ ≃ *λ*_*max*_, because growth of these organisms is far from equilibrium, which is included in the models through a high *K*_*eq*_ ≫ 1. In this regime, the C-source approaches its reservoir concentration while products and biomass approach zero due to wash out. Because of this behaviour, *X*_*growth*_ and the sEPR indeed increases monotonically in *D*, for both organisms. Close to the maximal dilution rate, they approach infinity.

**Figure 7.**
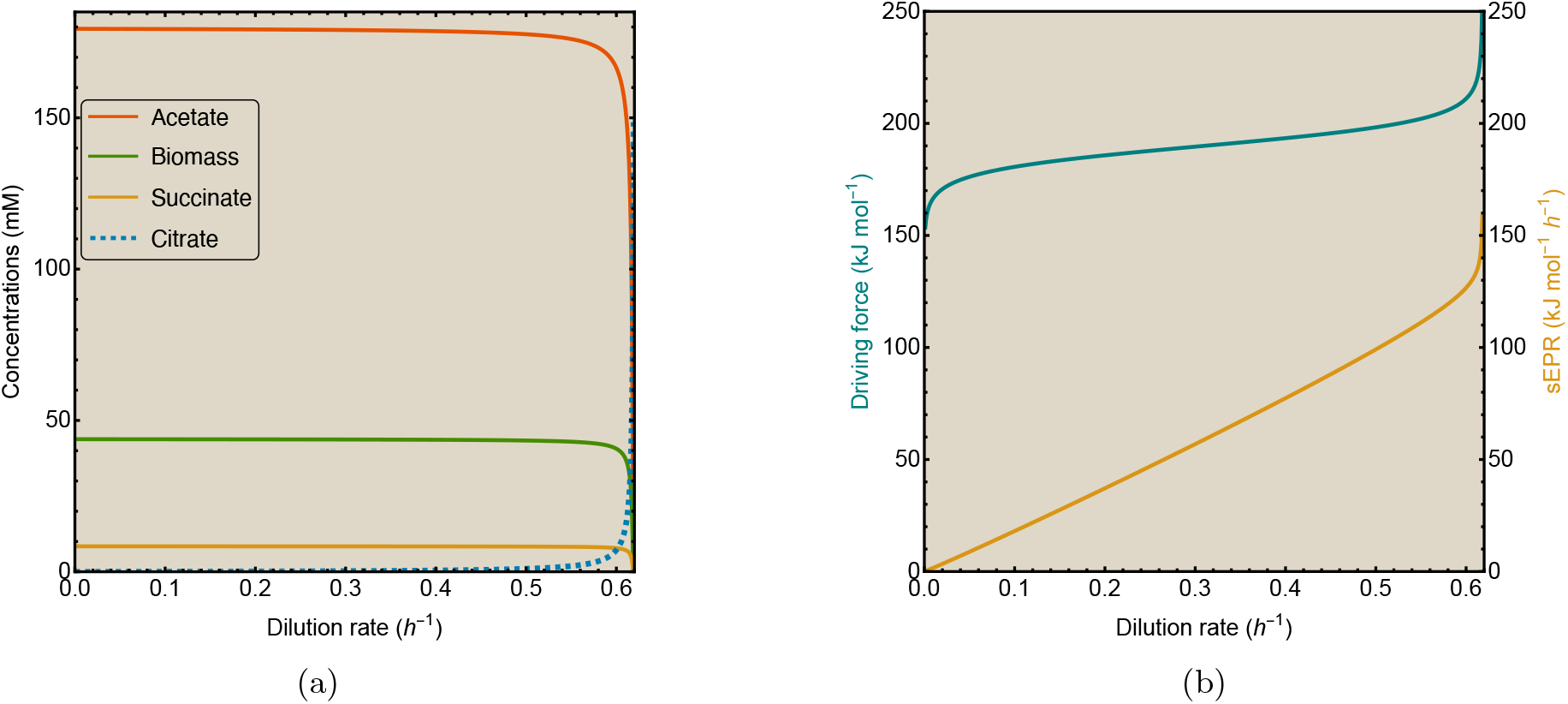
Results of the chemostat model for *K. aerogenes*. (a) Concentrations and (b) Thermodynamic quantities for anaerobic growth of *K. aerogenes* on citrate. Substrates are shown by dashed lines, products by solid lines. Model parameters are given in Table 1.

Mathematical modeling of the chemostat has a long history [Smith and Waltman, 1995, De Leenheer et al., 2006, Kuenen, 2019]. Our model assumes the chemostat contains a well-mixed liquid with only one microbial species, for which sticking to the walls of the bioreactor is negligible. Obviously, our models are not complete. Microbes invest some resources into maintenance, which decreases the observed yields [Pirt and Hinshelwood, 1965]. We do not think this has a large impact on our results. As wash out occurs for growth rates close to the maximal dilution rate *D*_*max*_, the model gives unrealistic results for both concentrations and thermodynamics in the chemostat close to *D*_*max*_; either they approach zero or infinity.

## Appendix E. sEPR always increases with growth rate for a fixed metabolic strategy in the chemostat

In general metabolic networks as considered in this work, the cell controls enzyme concentrations which determine the flux and the driving force. The example network in Section 3.3 illustrated that this can result in selection of a pathway with a higher specific flux but a lower driving force. When the macrochemical equation is fixed, this is not possible. We therefore claim that both the driving force and the sEPR rise with growth rate in chemostat conditions.

*Proof*. To see this, the dependence of the growth rate on the total driving force needs to be analysed. First, we study the steady-state solution of the chemostat model given by eq. (17). For any substrate *S*_*k*_, its steady-state concentration *s*_*k*_(*D*) decreases with increasing biomass concentration *b*(*D*). Similarly, for any product *P*_*l*_, its steady-state concentration *p*_*l*_(*D*) increases with *b*(*D*). Rewriting the dilution rate *D* in eq. (17) as function of *b* now gives that *D*^*′*^(*b*) *>* 0, as both the Monod relation and the contribution from thermodynamics decrease in *b*. Reversing this statement now yields that *b*^*′*^(*D*) *<* 0, which then implies that 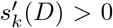 and 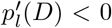. Inspecting then the subsequent relation (15) between the driving force and the dilution rate then shows that 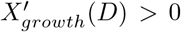. This shows that, for organisms in a chemostat modelled by a single fixed macrochemical equation, the driving force and therefore the sEPR rises with the growth rate. □

The relations between steady-state concentrations, driving force and the dilution rate are of course assumptions in the chemostat model, but they follow from observed behaviour and fundamental microbiology, such as the Monod relation. Furthermore, the growth rate in eq. (17) is exactly the same as the rate equation for mass-action kinetics in a chemical reaction network, except for the Monod term. So, the argument here follows a similar logic as for chemical reaction networks presented earlier, and therefore has the same conclusions.

## Appendix F. Modeling overflow metabolism with a mixing function

The models for overflowing *S. cerevisiae* are similar to the models presented previously. The only difference is that, after a critical dilution rate *D*_*c*_, the respirative and fermentative macrochemical equations are mixed.

This is due to the appearance of active intracellular constraints, which limits the microorganism in its behaviour and requires it to use other strategies to maintain its growth rate [de Groot et al., 2020]. So, constraints are now required to model these microbes accurately, which was not necessary before for the other organisms. Since these constraints do not appear naturally in our chemostat model, the mixing of strategies has to be included explicitly by introducing a mixing function *α*(*D*), which is defined as the fraction of resources invested in the fermentation strategy and is equivalent to a convex coefficient in an EFM decomposition of a steady-state flux vector (5). von Stockar and Birou [1989] have developed a similar method to model mixing of strategies in yeast.

Experimental data [Basan et al., 2015, van Hoek et al., 1998] suggests that uptake and excretion fluxes *q*(*D*) in a chemostat are affine in *D*, i.e., *q*(*D*) = *DY* (*D*) = *ν* + *τ D* for some *ν, τ* ≥ 0. Denoting the mixing function by *α*_*lin*_(*D*) for this case, the flux *q*(*D*) can be expressed as

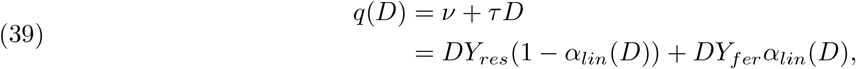

where the second form describes the change in the flux due a changing yield *Y* (*D*) as in eq. (21), which is affected by increasing investment in fermentation and decreasing investment in respiration. Solving this for *α*_*lin*_(*D*) yields

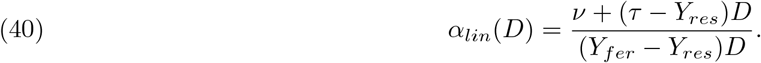

Parameter *D*_*c*_ is determined by the start of the metabolic shift, at which *α*_*lin*_(*D*_*c*_) = 0, which gives *ν* = (*Y*_*res*_ − *τ*)*D*_*c*_. At the ‘final’ dilution rate *D* = *D*_*f*_, yeast is using only fermentation, which translates into *α*(*D*_*f*_) = 1 implying *ν* = (*Y*_*fer*_ − *τ*)*D*_*f*_. Plugging in these two requirements gives the mixing function in the form (24)

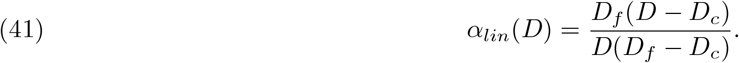

Note that *D*_*f*_ can not always be attained by the shifting organism. For example, yeast growing at maximal rate in aerobic batch conditions still uses some oxygen [Elsemman et al., 2022], so it seems to never reach the growth rate *λ* = *D*_*f*_.

To determine the parameters in the mixing function for each yeast model, it has to be fitted to data. Since the macrochemical equations from [Battley, 2013] for model 1 and [Heijnen and Dijken, 1992] for model 2 do not include accompanying chemostat data, the data from van van Hoek et al. [1998] is used. This contains flux data at different dilution rates for all important substrates and products of yeast. If the data and macrochemical equations are normalised to the same substrate (in this case glucose), the ethanol flux can be used to determine the fraction of resources invested in fermentation to achieve this flux at dilution rate *D*. This fraction should be equal to *α*_*lin*_(*D*). So, the mixing function can be fitted to this mixing data by determining these fractions for different dilution rates and thereby obtaining values for the parameters *D*_*c*_ and *D*_*f*_.

Performing these fits for our three yeast models can however result in a value for *D*_*c*_ *<* 0.28 h^−1^, while experiments show that *D*_*c*_ = 0.28 h^−1^. Furthermore, some fits also have a value for *D*_*f*_ *<* 0.4 h^−1^, while the data shows there is still some oxygen consumed at this dilution rate [van Hoek et al., 1998, Elsemman et al., 2022]. This would also imply that yeast has completely shifted to fermentation before growing at its maximal rate. Because of these potential issues, a different functional form for the mixing function is attempted. Based on the seemingly hyperbolic behaviour of the mixing data, we chose a hyperbolic form

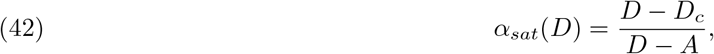

where *A* is a different fitting parameter. This function saturates slower to 1 than (41), as lim_*D*→∞_ *α*_*sat*_(*D*) = 1. Note that this function still satisfies *α*_*sat*_(*D*_*c*_) = 0. A disadvantage is that this choice results in nonlinear fluxes, while experimental data suggest these to be affine.

To choose either form (41) or (42), one would generally compare their goodness of fit to determine which form fits the data best. However, the data from [van Hoek et al., 1998] is so sparse that *R*^2^ values are extremely high (*>* 0.95). So, this is no reliable measure for comparison, as many different forms for *α*(*D*) with two fitting parameters could fit the obtained mixing data well. Therefore, we base the choice of the form for the mixing function on the values of the fitting parameters *D*_*c*_ and *D*_*f*_. We use *α*_*lin*_(*D*) as default, unless after fitting this to the mixing data we obtain a *D*_*c*_ *<* 0.28 h^−1^ or *D*_*f*_ *<* 0.4 h^−1^. In that case, we use *α*_*sat*_(*D*). Based on these criteria a mixing function is determined for each model, which are specified in Appendices G and H.

Note that the behaviour of the sEPR depends on the choice for the form of the mixing function. Just as the form (41) results in affine fluxes, it also results in an affine sEPR as function of *D*, while eq. (42) yields hyperbolic behaviour. So, the behaviour of the sEPR is therefore not uniquely determined, which makes it hard to draw strong conclusions. Nevertheless, we believe our models gives a description of the sEPR that is as accurate as possible for a theoretical analysis.

## Appendix G. Chemostat models for overflowing *S. cerevisiae* based on literature

Models 1 and 2 for overflowing yeast growing in a glucose-limited chemostat are based on macrochemical equations obtained from the literature. Model 1 uses macrochemical equations for respiration and fermentation of *S. cerevisiae* from [Battley, 2013], which are respectively given by

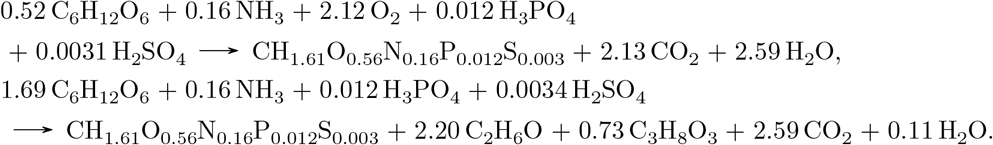

Based on the criteria as described in Appendix F, the mixing function used for this model for overflowing yeast is given by

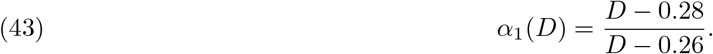

Model 2 uses macrochemical equations for respiration and fermentation of *S. cerevisiae* from Heijnen and Dijken [1992], which are respectively given by

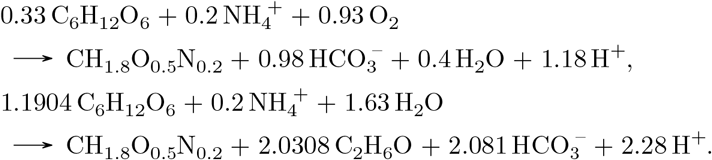

The mixing function used for this model for overflowing yeast is given by

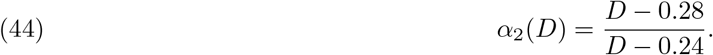

Both references use slightly different biomass compositions, and differ significantly in stoichiometric coefficients for glucose and ethanol (for fermentation). Also, oxygen and carbon-dioxide appear in some macrochemical equations, which are gases that are not completely dissolved in water. Therefore, their contribution to the concentration term in eq. (15) should only include the dissolved fraction. However, as all growth processes considered here are far from equilibrium, this effect on the total driving force for growth is negligible. It can therefore be assumed that oxygen and carbon-dioxide are completely dissolved in the bioreactor.

All other parameters of these models can be found in Table 1. The standard Gibbs energies are calculated using the thermodynamic tables given in the same reference as we obtained the corresponding the macrochemical equation from. It should be noted that Battley uses biological standard conditions, while Heijnen & van Dijken use non-biological standard conditions. Both use the thermodynamic reference frame.

## Appendix H. Chemostat model for overflowing *S. cerevisiae* based on data

Model 3 for overflowing yeast is directly based on the data from [van Hoek et al., 1998], whom measured uptake rates and effluxes of different metabolites during growth of *S. cerevisiae* in a glucose-limited chemostat. Macrochemical equations based on this data were reconstructed using the GEM Yeast8 [Lu et al., 2019]. Uptake rates and effluxes for all metabolites and trace elements were obtained with the GEM at different dilution rates 0 *< D <* 0.4 h^-1^, by lower bounds on ethanol and acetate efflux that correspond to the measured fluxes of these overflow metabolites at a certain dilution rate *D*, which was then also the lower bound of the growth rate. The constraint on the non-growth associated maintenance reaction was removed. Under these constraints the glucose uptake rate was minimised. This results in a list of uptake rates and effluxes for all constituents that satisfies elemental balance at each dilution rate. For overflowing yeast at *D > D*_*c*1_ these fluxes are a mix of different macrochemical equations (EFMs). An EFM enumerator [Terze, 2009] was used via CNApy [Thiele et al., 2021] to determine the EFMs that were used by the GEM at these dilution rates. For *D > D*_*c*1_ the GEM makes use of three EFMs to satisfy the three constraints for biomass, ethanol and acetate flux, which can be characterised as respiration, ethanol fermentation and acetate fermentation. The elemental composition of biomass was computed from the elemental balance condition. The GEM shows an inexplicable overconsumption of phosphate ions and overproduction of diphosphate, which seems to be a model incoherence and is therefore neglected. This results however in an oxygen unbalance, explaining the low coefficient for oxygen in the biomass composition.

The macrochemical equation for respiration, which is used both below and above *D*_*c*1_ is given by

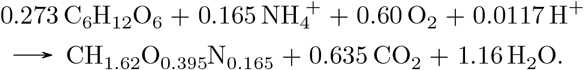

The macrochemical equation for ethanol fermentation is given by

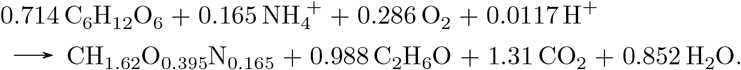

The macrochemical equation for acetate fermentation is given by

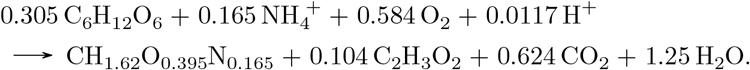

Note that all strategies make use of oxygen, which is energetically favorable and available in the medium of the GEM. Mixing coefficients, which describe the fraction of resources directed towards a strategy, were determined for each EFM by finding a unique flux and computing the fraction of glucose investment to attain this flux. Equivalently, we have computed the fraction of biomass synthesized by this EFM, as all EFMs produce biomass. This fraction is more convenient as the biomass yield is normalised at every *D*. The intuitive choice for the unique fluxes is the ethanol efflux for ethanol fermentation and acetate efflux for acetate fermentation. However, during the analysis it turned out that around *D* = 0.35 h^−1^, the sum of the mixing coefficients of these two fermentation strategies would exceed 1, which indicates that the cell would need to use more glucose than available to satisfy the constraints. As this is not possible, this behaviour hints at a second critical growth rate *D*_*c*2_ = 0.35 h^−1^, after which new strategies emerge with different yields that can satisfy the constraints. This is in line with other recent findings using genome scale modeling of yeast [Elsemman et al., 2022]. After this second shift at *D > D*_*c*2_, it turns out that both macrochemical equations for respiration and acetate fermentation change. The yields in the macrochemical equations change slightly and they start to produce ethanol as well. The ethanol fermentation strategy remains the same. This model therefore includes five macrochemical equations, three of which are given above. For *D > D*_*c*2_, respiration is given by

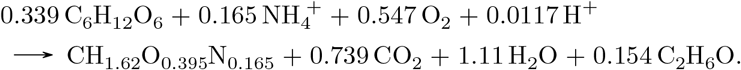

The macrochemical equation for acetate fermentation for *D > D*_*c*2_ is given by

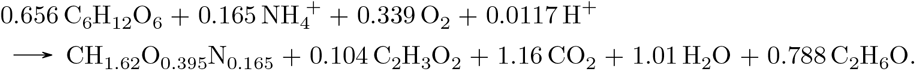

For these five macrochemical equations the corresponding standard Gibbs energies were computed using the table from Heijnen and Dijken [1992], as this was the most complete table for these reactants. To find the formation energy for biomass, its elemental composition was compared with an extensive list in [Popovic, 2019] containing formation energies for different microorganisms. The biomass sharing closest resemblance with our biomass composition has a formation energy of 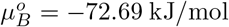. The resulting values are given in Table 1, next to other parameters for model 3. A control calculation with the table from [Heijnen and Kleerebezem, 2010] showed only small deviation of at most 3%, indicating that our method gives reasonable values for the standard Gibbs energies of macrochemical equations. It should be noted that the standard Gibbs energy for acetate fermentation is even more negative than for respiration, which contradicts the idea that respiration reduces glucose further and can therefore extract more free energy. However, looking at the macrochemical equations of respiration and acetate fermentation in both critical regimes, one can observe that acetate fermentation has more in common with respiration than actual (ethanol) fermentation. This can be seen from the high biomass yield on glucose, low carbondioxide efflux and low oxygen uptake rate. In general, the standard Gibbs energies for respiration for model 1 and 2 are much more negative than the value found for model 3. To investigate these two interesting features, more detailed data would be required than is currently provided by [van Hoek et al., 1998].

As there are two critical regimes, this requires four mixing functions (one for each fermentation strategy in each regime) to fit to the mixing coefficients. However, the data by [van Hoek et al., 1998] only includes 4 measurements for *D > D*_*c*1_ = 0.28 h^−1^, which is too sparse to properly fit different forms of the mixing function, as also explained in Appendix F. To make a better choice for the mixing function, the existing data is extended using a linear interpolation. With this interpolation the ethanol and acetate efflux for each 0.28 h^−1^ ≤ *D* ≤ 0.40 h^−1^ with a stepsize of 0.1 were determined and used as constraints in the GEM. Running the GEM now yields fluxes, macrochemical equations and thereby mixing coefficients at more values of the dilution rate than in the original data. It should be noted that this gave inconclusive results on the exact moment of the second shift, as multiple values for *D*_*c*2_ were compatible with a sum of mixing coefficients below 1. Based on [Elsemman et al., 2022] we have fixed the second shift at *D*_*c*2_ = 0.35 h^−1^. To summarise, the yields *Y*_*i/B*_(*D*) for this model of overflowing yeast in each growth regime are now given by

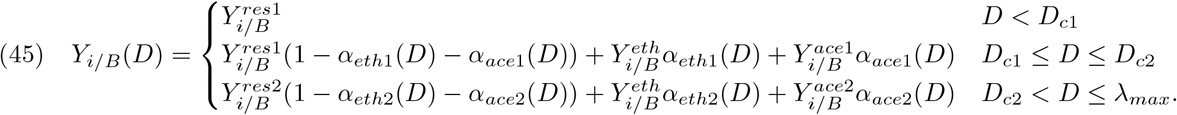

Similar equations can be written down for the (standard) Gibbs energies, just as in eq.(22). The four mixing functions for each fermentation strategy in each regime are given by

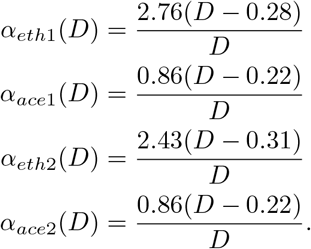

The mixing function for the acetate fermentation strategy is the same in both critical regimes. This is because only this EFM produces acetate, for which the efflux increases linearly in *D* according to the data. As this model contains two shifts and five EFMs, most fitting parameters in these four mixing functions lose their biological interpretation. Therefore, we use the default form (41) for all mixing functions in this model. Figure 4b shows the rates *q*_*i/B*_(*D*) = *DY*_*i/B*_(*D*) calculated with eq. (45) together with the data from van Hoek et al. for the most important reactants. This shows that model 3 describes the data reasonably well, except for the oxygen uptake rate. This is probably due to unrealistic behaviour of the GEM. The results of the chemostat model for this description of overflowing yeast in terms of five EFMs are depicted in Figures 5e, 5f.

## Appendix I. Derivation of quantitative criterion for behaviour of the sEPR

In Section 3.6 we have used a quantitative criterion involving an approximation of the sEPR to predict the behaviour of the sEPR for *D > D*_*c*_ for our three yeast models and other shifting organisms. Here we will present a derivation of this criterion.

For *D > D*_*c*_ the sEPR decreases if *ϕ*^*′*^(*D*) *<* 0. This derivative can be computed directly using the (mixed) yields and standard Gibbs energies (21). However, as Δ*μ*_*growth*_(*D*) is nonlinear and includes all substrates and products of the macrochemical equation, the result will not be insightful. Therefore, some assumptions are required to find a simple and clear criterion. The Gibbs energy dissipation can be written in terms of a standard Gibbs energy plus a concentration effect

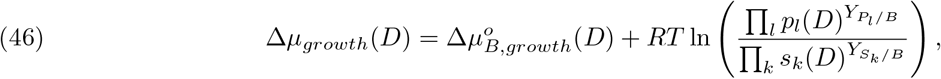

by applying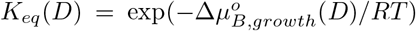. The dependence of 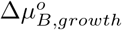 on the dilution rate follows from mixing of two metabolic strategies 1 and 2 as

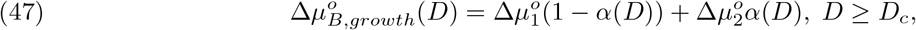

similar to eq. (22). Growth of all organisms considered in this work is far from thermodynamic equilibrium, as 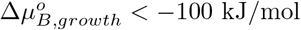 for all macrochemical equations. This means that the concentration effect is negligible for the Gibbs energy dissipation,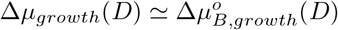. To derive a simple criterion, we assume that *α*(*D*) has the form (24) resulting in affine fluxes *q*_*j*_(*D*). Together, these assumptions result in an sEPR that is also approximately affine for *D* ≥ *D*_*c*_, as

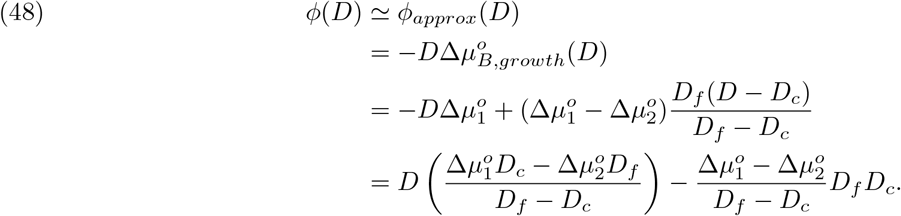

Hence, the sEPR decreases for *D > D*_*c*_ when

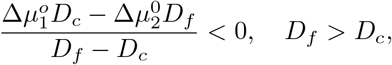

which may be summarised as

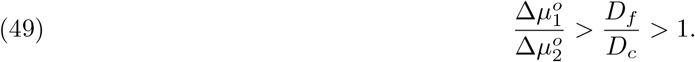

Each microbial species that displays a metabolic shift is characterised by a coordinate 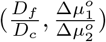. If this coordinate lies above the (dimensionless) diagonal given by the sEPR is predicted to decrease. 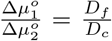 (the blue region in Figure 6), then the sEPR is predicted to decrease.

## Appendix J. Data for different organisms with a metabolic shift to predict their sEPR behaviour

This appendix contains information on growth and standard Gibbs energy data obtained for organisms that are represented in Figure 6. As briefly explained in Section 3.6, most of these organisms are represented by vertical lines. The reasons for this will be elaborated on here.

### J.1. Data for yeast models

For the three yeast models constructed in this work, coordinates 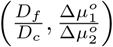 were determined and represented in Figure 6 as blue dots to predict the behaviour of their sEPR. For model 1 and 2 it should be noted that these use a mixing function of the form (42), which includes no parameter *D*_*f*_. In order to evaluate the criterion for these models, we compute *D*_*f*_ from mixing functions of the form (24) that were not used in the corresponding chemostat model, as otherwise the criterion can not be evaluated for these models. It should also be noted that *D*_*f*_ is ambiguously defined for organisms mixing more than two strategies, as happens for model 3. In that case we have calculated *D*_*f*_ as the dilution rate at which there would be only ethanol fermentation, as also the standard Gibbs energy of ethanol fermentation was used for 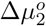.

Based on the coordinates found for the three yeast models, the prediction by the criterion is correct for models 1 and 3, but incorrect for model 2. Such a false prediction could follow from the approximation *ϕ*(*D*) ≃ *ϕ*_*approx*_(*D*). To check the reliability of this approximation, we have plotted *ϕ*(*D*) and *ϕ*_*approx*_(*D*) for each yeast model in Figure 8. One can conclude from this that *ϕ*_*approx*_(*D*) is indeed a crude approximation for models 1 and 3, but it does capture the qualitative behaviour of *ϕ*(*D*).

**Figure 8.**
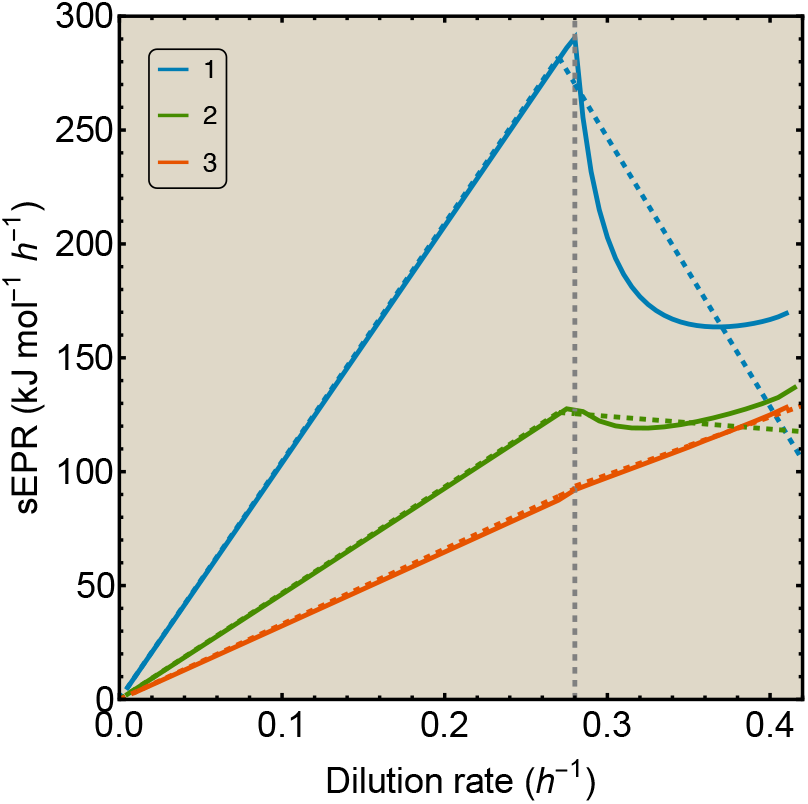
Approximation of the sEPR for the three chemostat models for overflowing yeast. Approximation is depicted as a dashed line. For model 1 and 3 it captures the characteristic behaviour, but not for model 2.

For model 2 the criterion predicted a decreasing sEPR, which is indeed the behaviour of *ϕ*_*approx*_(*D*). However, for this model it turns out that approximating the sEPR loses the key qualitative behaviour of *ϕ*(*D*), as this first shows a drop before rising again. As the criterion is independent of *D*, it can not predict such non-monotonic behaviour.

Nevertheless, we conclude that the criterion (49) is useful to predict behaviour of the (approximated) sEPR, as it depends only on a few model parameters. It therefore allows for a simple characterisation of thermodynamics of organisms with metabolic shifts.

### J.2. Data for other organisms with metabolic shifts

All organisms except *L. lactis, L. casei* and *S. ruminantium* show metabolic shifts in aerobic environments, from respiration at low growth rates to fermentation at high growth rates. Hence, these shifts are similar to overflow metabolism in yeast. The reason that these organisms are depicted by continuous vertical lines in Figure 6 instead of one point is because not all required data for evaluating the criterion (25) is available. For these organisms, data on critical growth rates *D*_*c*_ when fermentation is initiated is usually available. *D*_*f*_ is not measurable for most organisms, either because of chemostat limitations which decrease *D*_*max*_ *< D*_*f*_ or because, like yeast, the organism still respires a small fraction of its glucose uptake at maximal growth rate *λ*_*max*_. Instead, *λ*_*max*_ can be used as a lower bound on *D*_*f*_, as it is the maximal growth rate measured in batch at rich medium, which also gives a lower bound for *D*_*f*_ */D*_*c*_. This is a fair approximation to assess the criterion, as we are interested to find organisms that could potentially show a decreasing sEPR, which is more likely for lower *D*_*f*_. So, if *λ*_*max*_*/D*_*c*_ ≫ 1, this indicates a high probability of increasing sEPR, as the criterion for decreasing sEPR requires 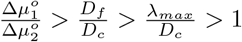.

A problem arises when examining the standard Gibbs energies of the two strategies that the organisms use during the shift. As we have observed for yeast, the standard energy 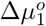 of respiration is usually available, but the standard energy 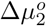 of the exact fermentation strategy used by the organism is harder to retrieve, for reasons explained in Appendix H. Therefore, the ratio 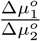 can not be calculated unambiguously for these organisms, which translates to an unknown *y*-coordinate in Figure 6 and consequently vertical lines for these aerobic metabolic shifts.

Table 2 contains the growth rate data for the organisms with metabolic shifts during aerobic growth, which determines their *x*-coordinate in Figure 6. For *S. kluyveri* [Møller et al., 2002] and *H. uvarum* [Venturin et al., 1995] it must be noted that even though the authors observed a metabolic shift, they still classified the yeast as being Crabtree-negative. Also the maximal growth rate of these yeasts is unknown, so an estimate for *λ*_*max*_ is obtained by extrapolating data for *S. kluyveri* and by taking the maximal experimental dilution rate for *H. uvarum. K. aerogenes* exhibits overflow metabolism in phosphate-limited chemostats [Neijssel and Tempest, 1976]. For *E. coli* W3110, *D*_*f*_ ≫ 0.8 h^−1^, as oxygen uptake seems to remain consant for this strain [Varma and Palsson, 1994]. For *E. coli* MG1655 the critical and maximal growth rates were measured by two different studies [Nanchen et al., 2006, Valgepea et al., 2010], finding different results, so both are included.

The organisms *L. lactis, L. casei* and *S. ruminantium* show metabolic shifts during anaerobic growth. Data for their critical growth rates *D*_*c*_ and *λ*_*max*_ are also contained in Table 2. *L. lactis* shifts from heterolactic fermentation at low dilution rates to homolactic fermentation at high dilution rates. The critical dilution rate is however ill-defined, as lactate is already produced at the onset of measuring [Goel et al., 2015]. A similar shift is observed in *L. casei* [De Vries et al., 1970] and *S. ruminantium* [Scheifinger, 1975]. These anaerobic organisms are also represented by vertical lines in Figure 6, but for a different reason than the aerobic organisms. This will be explained in the following appendix.

## Appendix K. General anaerobic catabolic shifts and methods to approximate Gibbs energy dissipations for growth

For the aforementioned and other anaerobic shifts, catabolic Gibbs energy dissipations are well-characterised for both strategies. These shifts also occur in the catabolic part of the growth process, but the (total) Gibbs energy dissipation is given by

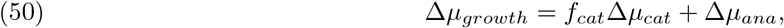

where *f*_*cat*_ represents the number of times the catabolic reaction runs to synthesize one mole biomass, if both processes are properly normalised. Hence, Δ*μ*_*growth*_ is also determined by anabolism and the coupling of these processes. As anabolism corresponds (mostly) to biosynthesis, it usually costs energy, so therefore Δ*μ*_*ana*_ ≥ 0 in most cases [Heijnen and Kleerebezem, 2010]. Due to coupling through ATP, anabolism is driven by catabolism. We have tried to circumvent these unknown parameters by comparing the ratios 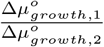 and 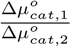 for two general metabolic strategies 1, 2. If the second ratio is larger than the first, we can use the criterion by substituting the ratio of the catabolic Gibbs energy potentials for the total Gibbs energy potentials of growth. There are indications that these ratios are comparable, for example the observed positive relation between the ATP yield and Δ*μ*_*cat*_ [Heijnen and Kleerebezem, 2010]. However, because no strict bounds exist on both Δ*μ*_*ana*_ and *f*_*cat*_, the required inequality can not be proven.

So, in order to compute the Gibbs energy potentials of both strategies that are exploited during the anaerobic shift, which are required to evaluate the criterion (49), the values for Δ*μ*_*ana*_ and *f*_*cat*_ need to be calculated for an organism exploiting such a shift. In principle, it is possible to estimate an anabolic reaction and the corresponding Gibbs energy change. Heijnen and Kleerebezem [2010] and Ebenhöh et al. [2023] describe similar methods to obtain a general anabolic reaction that only depends on the carbon source, biomass composition and their degree of reduction. Kleerebezem and van Loosdrecht [2010] use redox half reactions to get a better estimate for the anabolic reaction. However, both methods assume that every organism with similar biomass composition growing on glucose can be represented by the same anabolic reaction. The factor *f*_*cat*_ is directly related to the yield of biomass on the carbon source 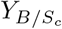 [Kleerebezem and van Loosdrecht, 2010]. This yield is usually experimentally determined for a single growth strategy.

However, when two strategies are mixed for *D > D*_*c*_, also the yield is a function of the growth rate. It is therefore hard to determine 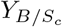 for the second strategy, without constructing a complete chemostat model and determining a mixing function for such an organism. This implies that *f*_*cat*,2_ for strategy 2 can not be determined in most cases, or can only be estimated roughly.

In conclusion, the methods mentioned here can only give crude estimations of 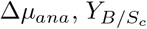 and *f*_*cat*_, and hence also a crude estimation of the complete macrochemical equation and its Gibbs energy potential. During this study it was observed that yields and (standard) Gibbs energy potentials in the macrochemical equations have a large impact on the behaviour of the sEPR. Therefore, we consider a prediction for anaerobic organisms based on the criterion (49) and these crude estimations to be unreliable. Therefore, as thermodynamic data is incomplete, also these anaerobic shifts are depicted as vertical lines in Figure 6.

## Footnotes

1 In the literature formation energies are often presented as 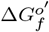, but as we only look at thermodynamic quantities per mole of a compound, we consistently denote this as 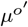^*′*^.

2 We hope not to confuse the reader with the twofold use of the subscript _*B*_, which denotes the biomass for the yields *Y*_*i/B*_, but biological standard conditions for the standard Gibbs energy potentials 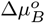.

3 This extra assumption is now required; since the conversion can differ per EFM, *v*_*r*_ is not necessarily nonzero in every EFM.

